# Repeated genetic adaptation to high altitude in two tropical butterflies

**DOI:** 10.1101/2021.11.30.470630

**Authors:** Gabriela Montejo-Kovacevich, Joana I. Meier, Caroline N. Bacquet, Ian A. Warren, Yingguang Frank Chan, Marek Kucka, Camilo Salazar, Nicol Rueda, Stephen H. Montgomery, W. Owen McMillan, Krzysztof M. Kozak, Nicola J. Nadeau, Simon Martin, Chris D. Jiggins

## Abstract

Repeated evolution can provide insight into the mechanisms that facilitate adaptation to novel or changing environments. Here we study adaptation to high altitude in two divergent tropical butterflies, *H. erato* and *H. melpomene*, which have repeatedly and independently adapted to high elevations on either side of the Andean mountains. We sequenced 518 whole genomes from elevational transects and found many regions under selection at high altitude, with repeated genetic differentiation across multiple replicates, including allopatric comparisons. In contrast, there is little ‘molecular parallelism’ between *H. erato* and *H. melpomene*. With a further 85 whole genomes of five close relatives, we find that a large proportion divergent regions have arisen from standing variation and putative adaptive introgression from high-altitude specialist species. Taken together our study supports a key role of standing genetic variation and gene flow from pre-adapted species in promoting parallel genetic local adaptation to the environment.

## Introduction

Understanding how organisms adapt to the environment is a central goal of evolutionary biology and highly relevant given the pace of global change. One approach is to explore the repeatability of local adaptation in the wild in order to understand whether phenotypic and genetic changes are predictable. On the one hand, repeated adaptation to similar environments can act as a ‘natural experiment’ and provide the means to identify the targets of selection, by distinguishing locally adaptive from neutral or globally beneficial changes^1^. On the other hand, these scenarios can allow us to test whether the same loci are repeatedly targeted across populations and species^2^. Despite many studies reporting repeated adaptation involving the same genes or alleles across lineages^3–5^, which we here term ‘molecular parallelism’, we know relatively little about the evolutionary mechanisms that facilitate it.

Three main mechanisms can give rise to molecular parallelism in repeated adaptation (Fig.1). Genetic variation upon which selection repeatedly acts may arise via independent mutations at the same gene or locus^6^. Beneficial variants may be recruited from ancestral standing variation^7^ or shared across populations of the same species via migration and gene flow^8^. Lastly, gene flow between species can facilitate the introgression of adaptive alleles^9–11^. A combination of these mechanisms may also be at play, for instance the high altitude adaptation Tibetan-EPAS1 haplotype was introgressed from Denisovan hominins but remained as neutral standing variation before positive selection occurred^12^.

**Figure 1.**
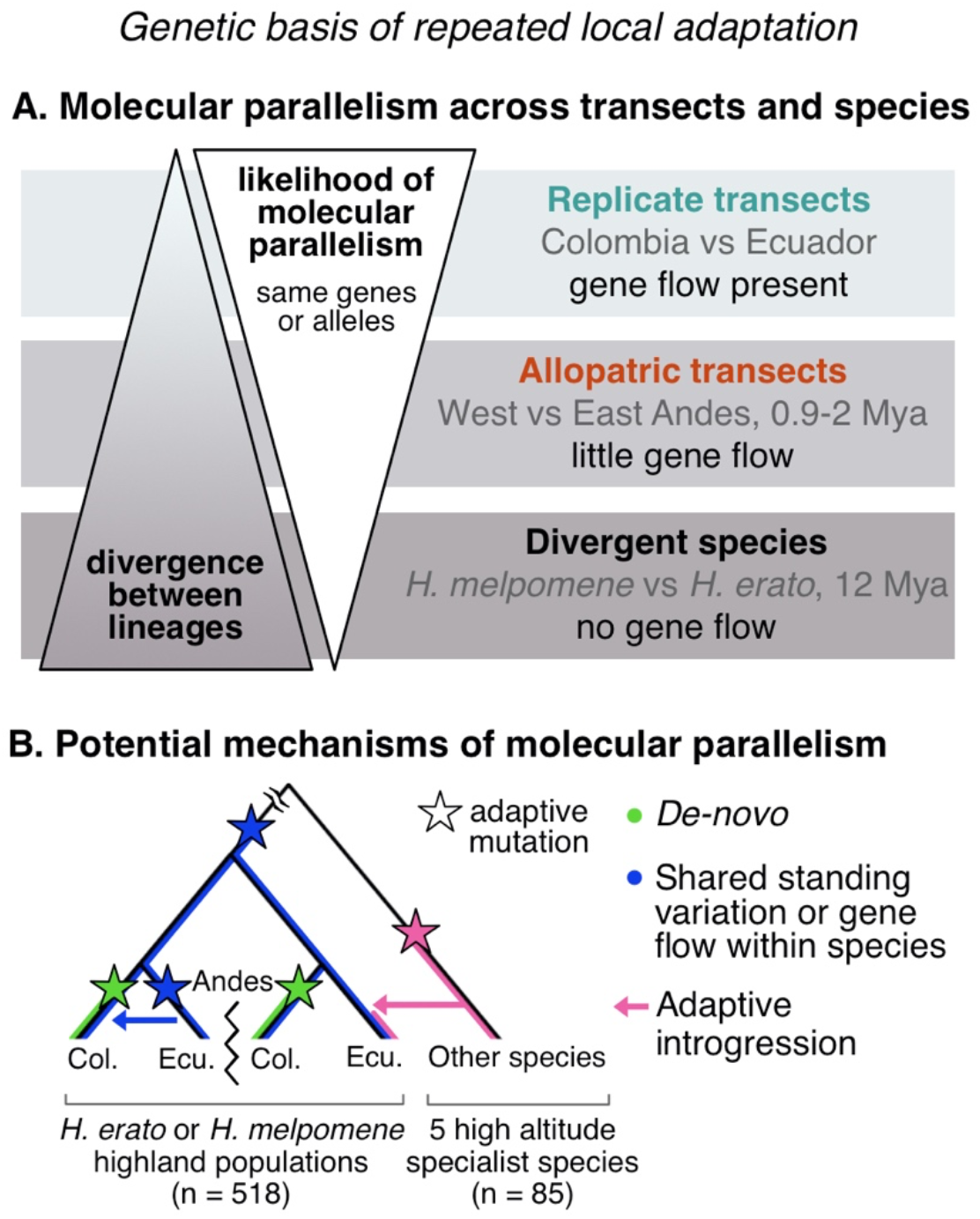
The study of repeated adaptation to the environment and the mechanisms potentially facilitating it. A) We hypothesise that increasing divergence between the lineages under study reduces the likelihood of molecular parallelism (same genes or alleles) underlying repeated adaptation to the environment. In this study, we test this hypothesis by sampling replicate (within sides of the Andes) or allopatric (across sides of the Andes) altitudinal transects of the same species, i.e. connected via gene flow or not (divergence times indicated in Million years ago, Mya), and all replicated in two divergent *Heliconius* species. B) Three main mechanisms can give rise to the genetic variation upon which selection acts repeatedly, giving rise to molecular parallelism: (i) adaptive *de-novo* mutations independently arise in two or more lineages, (ii) existing shared standing variation is repeatedly selected across lineages or shared via gene flow within species, and (iii) adaptive alleles are shared via gene flow across species (adaptive introgression). We tested for the relative importance of these mechanisms with a range of analyses on the relevant transect comparisons. Illustrative tree including four highland populations from four transects (with the Andes preventing gene flow between replicate transects across sides) of our focal species, either *H. erato* or *H. melpomene*, and a lineage of a related high-altitude specialist species from which adaptive introgression is plausible. Sample sizes for the full datasets (including lowland populations when present) are shown in brackets.

The likelihood of molecular parallelism and the relative importance of each mechanism may be largely dependent on the divergence between the lineages under study (Fig. 1), but this has seldom been empirically tested^3^. For instance, populations that diverged recently and retain a large pool of standing genetic variation tend to reuse preexisting alleles during repeated adaptation, as seen in freshwater adaptation in sticklebacks^13^, crypsis in beach mice^14^, or coastal ecotypes of bottlenose dolphins^7^. Similarly, organisms that readily hybridise in the wild are more likely to share beneficial alleles via adaptive introgression. This allows populations to rapidly adapt to, for instance, novel anthropogenic stressors such as pollutants^15^ or insecticides^16^.

Anthropogenic change is forcing organisms to move, adapt, or die, with many predicted to expand their ranges towards the highlands to escape warming and degrading lowland habitats^17^. Despite insects making up about half of all species that have ever been described^18^, we know very little about the genomics or predictability of adaptation to altitude (but see^19,20^), especially in the tropics. The type and genomic architecture of the trait under study may also determine its predictability^21^. Phenotypes controlled by few, large-effect loci typically show predictable genetic paths of evolution, such as melanic colouration in mammals, fish, and birds^22–25^, perhaps due to selective constraints on genetic pathways^21,26,27^. Organisms adapting to complex environmental challenges that face multifarious selective regimes may show less predictable patterns, with functional redundancy between genes allowing for different combinations of alleles to achieve similar phenotypes^28,29^. Thus, understanding the relative importance of these mechanisms in determining the predictability of adaptation to the environment could inform future research into conservation strategies to protect biodiversity^30^.

Here, we study the genetic basis of repeated adaptation to high altitude in two divergent tropical butterflies, *H. erato* and *H. melpomene*. These aposematic, toxic species have very wide ranges and co-mimic each other across South America, commonly found from sea level to around 1600 m^31^. In contrast, most other species in this genus have specialised to highland Andean or lowland Amazonian habitats, with topography and climate shown to correlate with speciation rates across the clade^32^. Phenotypic differences between highland and lowland populations of *H. erato* and *H. melpomene* have recently been identified, such as in wig shape^33,34^ or heat tolerance^35^. We search for signatures of local adaptation to high altitude with extensive sampling that harnesses the power of natural spatial replication within and across sides of the Andes, in order to assess the extent of molecular parallelism in adaptation to high altitude. We quantify parallelism at multiple levels of divergence: (i) replicate transects within sides of the Andes connected via gene flow, (ii) trans-Andean allopatric transects with no gene flow at these latitudes, and (iii) two species that diverged 12 million years ago (Fig. 1A). Furthermore, we test whether the same haplotypes are under selection across transects and search for the mutational origin of candidate adaptive alleles. Overall, this large empirical study deepens our understanding of how organisms adapt to new environments and identifies both standing genetic variation and adaptive introgression from pre-adapted species as important mechanisms facilitating local adaptation.

## Results and Discussion

### Divergence and diversity across elevations and transects

To study adaptation to altitude in *H. erato* and *H. melpomene*, we used whole-genome data from 518 re-sequenced individuals, 444 of which were newly sequenced for this study. Samples were collected from 111 different locations, which we grouped into 30 populations, corresponding to four transects: Colombia West/East, and Ecuador West/East (Fig. 2, Table S1). In each transect, populations were either in the highlands (~1200 m) or lowlands (~200m), itself divided into nearby or distant lowland sites (Fig. 2 A). The Andes act as a barrier to gene flow at these latitudes, with populations on opposite sides of the Andes thought to have split ~0.9 and ~2 million years ago for *H. erato* and *H. melpomene*, respectively^36–38^. Individuals of each species clustered strongly into Western and Eastern groups in genome wide PCAs (S.I., Fig. S1). In PCAs that only included populations from replicate transects of the same side of the Andes (two per species), structuring by altitude was absent in all but one comparison, *H. erato* East, where the *H. erato* highland population in Colombia corresponds to a different colour pattern subspecies and diverged moderately from other populations (S.I., Fig. S1). Intraspecific pairwise differentiation between populations on the same side of the Andes increased with geographic distance but was generally low (F_st_ <0.1; S.I., Fig. S2). The effective replication over space and extensive gene flow within transects provide a powerful setting to study the genomics of parallel adaptation to altitude in the wild.

**Figure 2.**
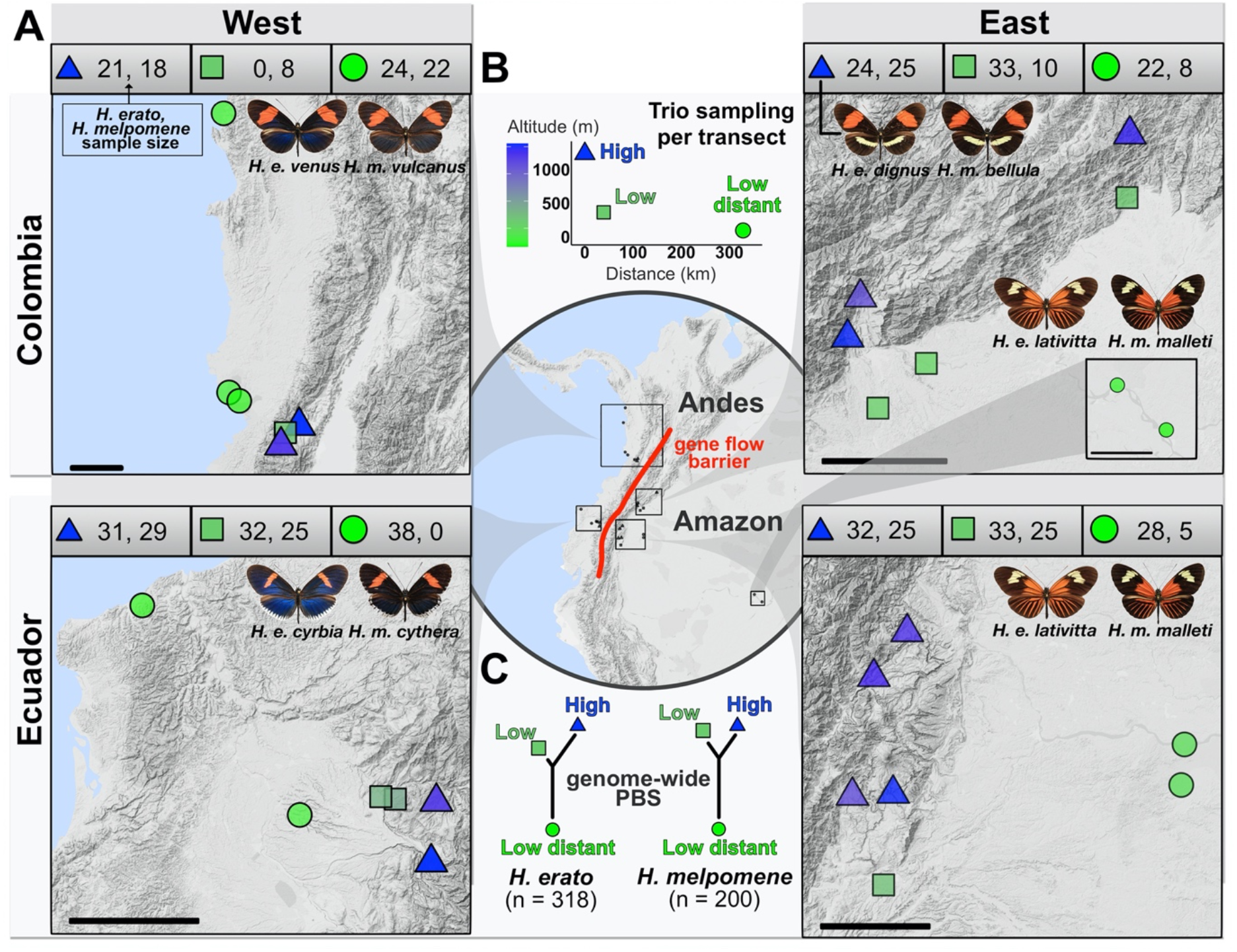
Sampling design. A) Elevation map of the 30 populations sampled for this study in four geographical transects (Colombia West/East, Ecuador West/East). More details of each population can be found in Table S1, number of whole-genome sequences included per elevation and transect is indicated above each map (*H. erato, H. melpomene*). The black scale bar represents 25km. *Heliconius* subspecies present are depicted per transect (credit: McGuire Center for Lepidoptera and Biodiversity, Florida Museum of Natural History), note that only Colombia East has different subspecies in the highlands compared to the lowlands. B) Plot depicts trio sampling scheme, with mean altitudinal and geographical distance of the three population types (high, low, low distant) for both species. C) Mean genome-wide Population Branch Statistic (PBS) trees averaged across the four transects per species and respective total sample size in brackets.

Genetic distance between populations can be increased by environmental differences that affect dispersal or survival of locally adapted migrants^39,40^. We tested for such isolation by environment, in our case altitude, with pairwise F_st_ across all populations of the same side of the Andes and species. At similar geographic distances, genetic differentiation was higher when comparing highland vs. lowland populations than when comparing lowland vs lowland populations. Isolation by altitude could be driven by local adaptation reducing gene flow between elevations or due to many adaptive sites diverging across the genome. This difference was stronger in *H. erato* than in *H. melpomene* (S.I., Fig. S2). F_st_ was generally highest when comparing two highland populations at short distances, despite using a topographically informed ‘least-cost path’ as our measure of geographic distance (S.I. Fig. S3). This could indicate topographical barriers decreasing gene flow, beyond what was captured by the least-cost path, or local adaptation in the highlands leading to increased selection against migrants. A pattern of isolation by environment could also arise due to, for instance, different demographic histories and higher levels of inbreeding in range-edge populations^41–43^. We found no consistent differences in nucleotide diversity (π) between elevations (S.I., Fig. S4). Tajima’s D was negative across populations, characteristic of population expansion, but generally less negative in highland populations, suggesting less pronounced expansion or more recent/ongoing contractions in the highlands (S.I., Fig. S4). Thus, both heterogenous demographic histories and selection against locally adapted migrants across elevations may lead to genome-wide isolation by environment.

### Parallel high-altitude differentiation detected with Population Branch Statistics

To identify genomic regions with high-altitude specific differentiation we calculated Population Branch Statistics (PBS) for three transects in each species, and F_st_ for the remaining two transects with more limited sampling (*H. erato* Colombia West, *H. melpomene* Ecuador West; Fig. 2A). PBS was originally developed to study high-altitude adaptation in humans^44^ and can distinguish between global and lineage specific differentiation by constructing a trifurcating population tree based on F_st_ that includes a geographically distant population^15,42,45–49^. By attributing fractions of differentiation to each branch, PBS identifies genomic regions disproportionally diverged in the focal population, consistent with positive selection or relaxed constraint^44,50^.

When assessing genome-wide average PBS trees the longest PBS branches corresponded to the low distant populations of both species, as expected under a model of neutral isolation by distance (Fig. 2B). *H. erato* had a consistently longer high-altitude branch compared to the lowland, but geographically nearby population, which could be indicative of increased drift or extensive local adaptation to altitude throughout the genome. Indeed, we detected many regions across the genome strongly differentiated in high altitude populations across transects and species (Fig. 3). We defined High Differentiation Regions (“HDRs” hereafter) by adding a ±50 kb buffer around outlier windows, i.e. those with zPBS_high_ (high-altitude branch) or zF_st_ values above 4 (standardised Z-transformed, equivalent to > 4 SD), and merging overlapping intervals into discrete regions. The transects for which only two populations were sampled (F_st_), had a higher number of HDRs: 400 and 405 HDRs, covering 11.4% and 17.1% of the genome for *H. erato* and *H. melpomene*, respectively (compared to, on average, 229 PBS-based HDRs covering 8%; details on S.I., Note S1). This likely reflects the property of PBS to discern between population-specific and globally differentiated alleles.

**Figure 3.**
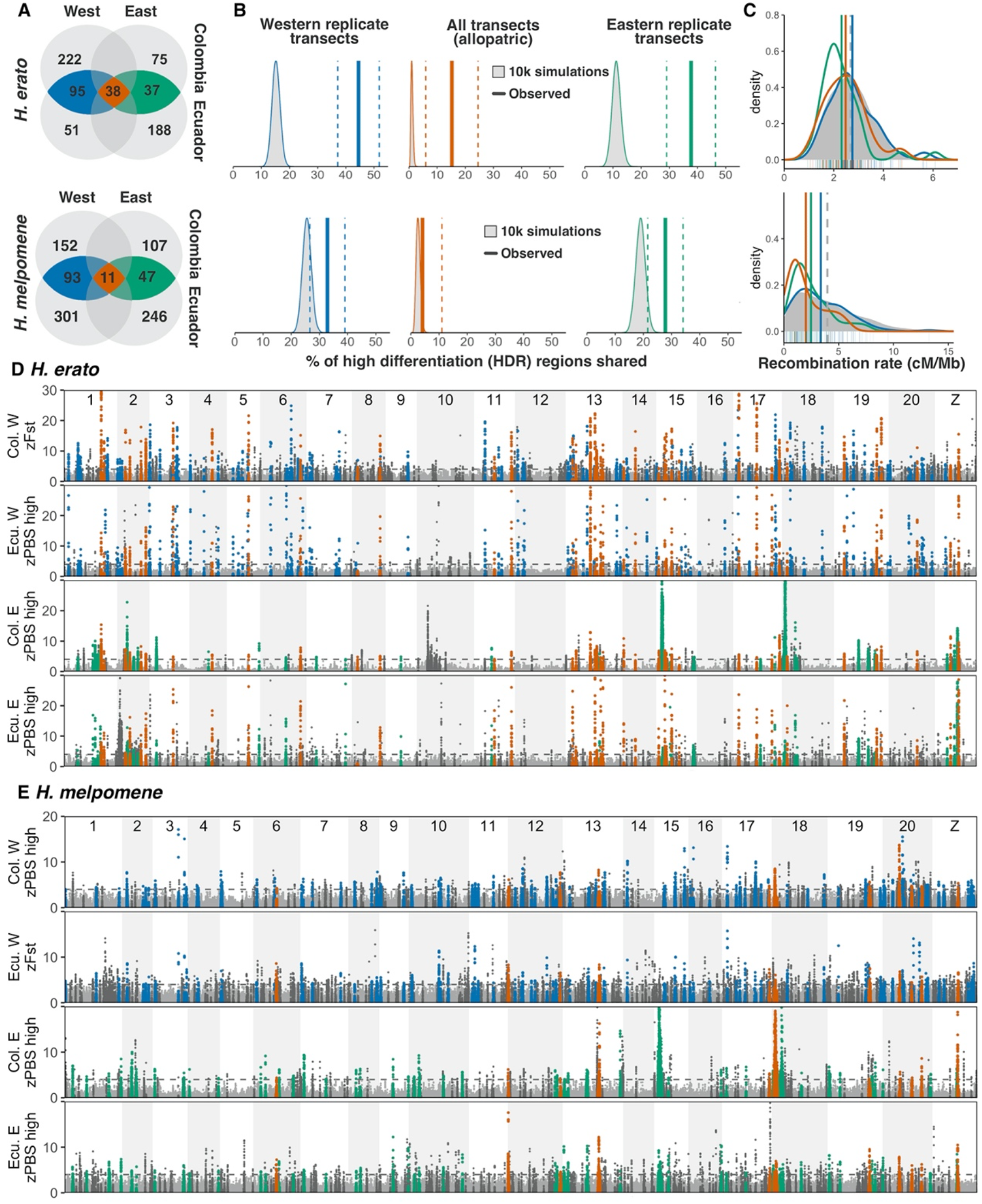
Molecular parallelism in PBS/F_st_ regions of differentiation across eight altitude transects of *H. erato* and *H. melpomene*. A) Number of high-differentiation regions per species (HDRs, including ± 50kb buffers), in blue/green if shared across replicate transects within sides of the Andes (SHDR: blue=within West, green=within East) and in red those additionally shared across allopatric transects, i.e. shared across all four transects (also SHDR). B) Vertical lines represent percentage of outlier windows shared across transects (jackknife resampling confidence intervals as dashed lines), compared to 10,000 simulations (grey distributions). C) Comparison of SHDR (coloured) and genome-wide (grey) recombination rates (cM/Mb). D-E Patterns of highland-specific differentiation (zPBS_high_) across the genome in four transects of *H. erato* (D) and *H. melpomene* (E). In the two transects where only two populations were sampled zF_st_ is presented. Horizontal dashed line indicates threshold of 4 standard deviations from the mean. HDRs private to one transect are highlighted in dark grey.

To test for molecular parallelism in local adaptation, we assessed whether the same individual HDRs were repeatedly found at high altitude across replicate (same side of the Andes) or allopatric (opposite side of the Andes) transects within each species (Fig. 1A). In *H. erato*, 45% (±3.8 SD) and 38% (±4.4 SD) of HDRs overlapped between replicate transects within the Western and the Eastern Andes, respectively (shared HDRs, SHDRs hereafter; Fig. 3D blue, green). Of those SHDRs, more than a third were also shared across allopatric transects not connected via gene flow, 15% of the total HDRs (allopatric SHDRs hereafter; Fig. 3D red). *H. melpomene* had a slightly lower percentage of HDRs shared within sides of the Andes (West 33% ±11 SD; East 27% ±11 SD), but very few shared across sides of the Andes (allopatric SHDRs: 4% of the total). We then tested if the observed level of sharing was higher than expected under a null distribution of genomic regions, obtained by assessing sharing in randomly placed blocks across the genome of the same size and number as observed HDRs per transect. *H. erato* HDR sharing was higher than predicted by our simulations in all three comparisons (replicate Eastern/Western and all transects, Fig. 3B), whereas the low levels of allopatric parallelism in *H. melpomene* did not differ from the null distribution (Fig. 3B, red). The difference between the observed and simulated extent of molecular parallelism of HDRs was robust to the buffer size around outlier windows used for assessing overlap (S.I. Note S1). Our selected buffer of ±50kb is conservative given that well-studied regulatory regions of colour pattern loci in these species can be up to 100kb from the coding sequence itself^51^. On average, 15.6% of SHDRs detected overlapped between species, but this fell within the simulated overlap levels given the number and size of SHDRs in each species (mean overlap = 14.06 % ±1.19).

Together, we show that there are high levels of parallelism in high-altitude differentiation within species, but little between the two species. On the one hand, molecular parallelism within species could be facilitated by the high levels of nucleotide diversity observed, indicating a large pool of shared variants upon which selection can repeatedly act. Furthermore, gene flow between populations across replicate transects (within sides of the Andes) could facilitate the recruitment of new or standing adaptive alleles, as expected from theory^52^ and seen in other systems such as maize, *Arabidopsis*, or sticklebacks^3,8,53^. On the other hand, the lack of significant molecular parallelism in altitude candidate loci between *H. erato* and *H. melpomene*, whose clades diverged 12 million years ago^38^, is in stark contrast with colour patterns^54^, where near-perfect local Müllerian mimics have arisen repeatedly in both species through independent mutations at a handful of conserved loci^51,55^. This difference in the extent of molecular parallelism might be explained by the nature of the trait under study: adaptation to altitude is multifarious and, as shown here, involves many genes. Genetic redundancy in polygenic adaptation may lead to evolution taking different paths to reach similar solutions, as shown for quantitative iridescence colouration in these two *Heliconius* species^56^ or in recent experimental evolution of thermal adaptation in *Drosophila*^28,57^. In contrast, the major effect loci that underlie switches in discrete colour patterning may favour genetic predictability even across divergent taxa, especially when selection is strong and reaching adaptive peaks requires large phenotypic shifts.

### Highly differentiated regions show additional signatures of selection and, generally, are not associated with low recombination rates

We tested whether highly differentiated genomic regions shared across transects (SHDRs) showed additional evidence of positive selection by computing difference in nucleotide diversity (π) across elevations (Δπ =π_high_-π_low_), deviation from neutrality in site frequency distributions (Tajima’s D), and absolute divergence (D_xy_) for the same 5 kb windows. Processes other than positive selection, such as background selection, can decrease within-population diversity and thus lead to increased relative differentiation (F_st_), especially in the absence of gene flow between populations^58,59^. Thus, it is important to test for enrichment of different selection statistics to strengthen our inference of locally adaptive loci. A reduced Δπ compared to the background would indicate that a selective sweep in the highlands reduced nucleotide diversity compared to the lowlands. In selective sweep regions, Tajima’s D is expected to be low, as regions with selected haplotypes that rapidly increased in frequency would have an excess of rare alleles. Finally, absolute sequence divergence (D_xy_), is expected to be high in old selective sweeps or variants, and less affected by genetic variation within populations than relative measures of differentiation such as F_st_^60^.

SHDRs were considered outliers for Δπ, Tajima’s D, or D_xy_, if the observed maximum or minimum values within SHDRs <10^th^ percentile (or >90^th^ in Tajima’s D) of the simulated values, obtained from 10000 permutations that randomly placed blocks of equal number and size to observed HDRs across the genome. Of the SHDRs differentiated on both Ecuadorian and Colombian transects but in one side of the Andes only, i.e. across replicate transects, on average 74% and 48% of *H. erato* and *H. melpomene* SHDRs, respectively, were at least outliers for one other statistic, in addition to zPBS/zF_st_ (Fig. 2 A grey). Of the *H. erato* and *H. melpomene* SHDRs shared across all transects (allopatric SHDRs), 94% and 86% had at least one additional outlier statistic, respectively (Fig. 4A, C). In *H. erato*, SHDRs were often outliers for both, high D_xy_ and reduced Tajima’s D (36% of SHDRs with additional outlier statistics, on average). In contrast, *H. melpomene* SHDRs were rarely outliers for Tajima’s D, whereas 22-33% of them were outliers for both Δπ and D_xy_. This could point towards different selection histories in each species, with *H. erato* showing signatures of recent or ongoing selective sweeps leading to an excess of rare alleles (negative Tajima’s D). The Andean split is dated ~0.5-1 million years older in *H. melpomene* populations^36,37^ and their altitudinal range is wider than that of *H. erato*, which is rarely found >1500 m at these latitudes. Thus, it is possible that *H. melpomene* SHDRs represent more ancient sweeps, reflected in the high prevalence of SHDRs outliers for D_xy_. Alternatively, high absolute sequence divergence (D_xy_) between elevations could be indicative of selected haplotypes arising through adaptive introgression from other species into the highland populations.

**Figure 4.**
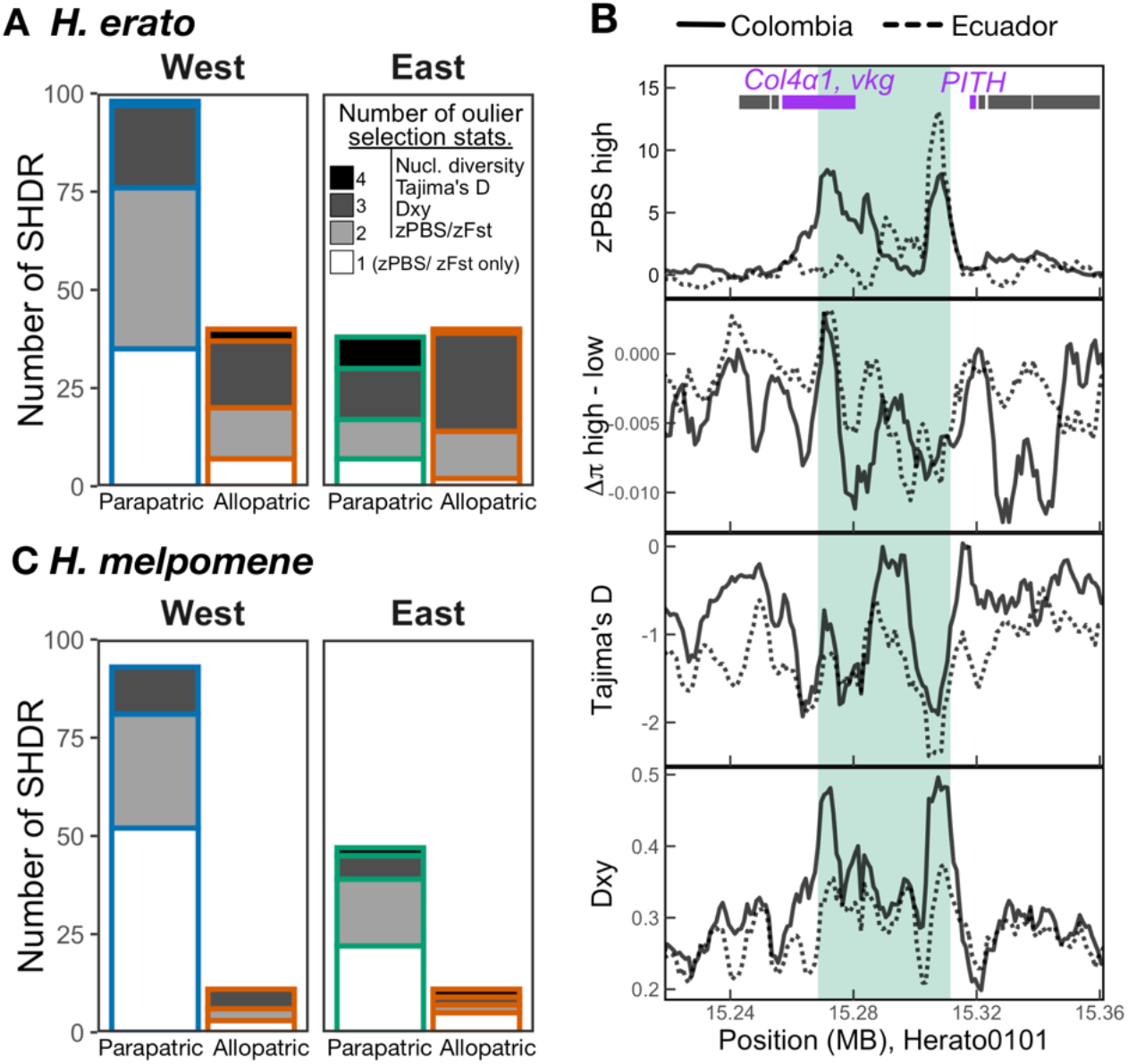
Signatures of positive selection across shared high differentiation regions (SHDRs). Number of SHDR with additional outlier selection statistics in *H. erato* (A) and *H. melpomene* (C), statistics included were nucleotide diversity difference between highlands and lowlands (Δπ), Tajima’s D, and absolute genetic differentiation (Dxy). Shading indicates number of statistics that were above 90^th^ percentile of simulations, white=1 (only zPBS), light grey=2, dark grey=3, and black=4 statistics. Example close-ups of regional zPBS highland values and statistic patterns in Eastern SHDR (#005, B). Each line represents the values for one of the two Eastern transects, solid lines are transects in Colombia and dashed in Ecuador. In this example, all three additional selection statistics ranked as outliers among simulations. Green shading highlights the region of the eastern SHDR with zPBS_high_ > 4.

As an additional independent line of evidence that SHDRs are under selection, we checked for overlaps with selection statistics and altitude-associated regions obtained from an altitudinal transect in southern Ecuador on the East of the Andes, sequenced with a new linked-read technology called ‘haplotagging’ (Meier *et al*., 2021). We found that, on average, 59% (± 10 SD) of Eastern SHDRs of each species overlapped with at least one additional outlier selection statistic estimated with the haplotagging dataset, whereas, as expected, fewer Western SHDRs did (32% ±8 SD, on average; S.I., Note S2, Fig. S5). In contrast, SHDRs shared in all transects showed high levels of overlap with haplotagging-derived selection statistics in all transects (52% ±15 SD, on average).

We tested if SHDRs were associated with low-recombining regions. Regions of high differentiation and low recombination could be indicative of purifying selection against deleterious mutations (background selection) or maladaptive introgression^59,61,62^. Background selection has been shown to be a major driver of differentiation landscapes between populations with little gene flow^60^ but is less plausible when populations readily exchange genetic material, as in this study^63,64^. However, in low recombining regions, selection may be more efficient due to reduced effective gene flow and segregation of coadapted alleles, and thus facilitate the maintenance of locally adaptive loci^59,65,66^. Several strongly selected *Heliconius* colour pattern loci have been previously associated with regions of low recombination (Fig. 3 chromosomes 15 and 18)^55,67^. Nevertheless, here we found that recombination rate at SHDRs did not differ from background levels, except in comparisons that included strongly selected colour pattern loci (Fig. 3C, S.I. Fig. S6). Overall, these additional signatures of selection strongly support the action of repeated divergent selection at high altitude rather than background selection driving the differentiation detected at SHDRs.

### Known genes of interest overlap with SHDRs

We retrieved 908 and 747 genes overlapping with SHDRs in *H. erato* and *H. melpomene*, respectively. With so many potential targets of selection within SHDRs, we do not attempt to infer biological function or adaptive significance from the whole gene set. Instead, we checked for overlaps with regions recently associated with wing shape variation across an altitudinal cline of Southern Ecuador in *H. erato* and *H. melpomene*^34^. Rounder wings are generally associated with high altitude across 13 species of *Heliconius*^33^ and subtle altitude-associated wing shape variation is highly heritable in *H. erato* and *H. melpomene*^34^. We found that five out of 12 previously identified candidate wing shape loci^34^, overlapped with SHDRs in *H. erato*, three of which corresponded to SHDRs detected in all transects. In contrast, only two wing shape loci (out of 16) overlapped with *H. melpomene* SHDRs, one of which was an allopatric SHDR. The number of overlaps between candidate wing shape loci and SHDRs in *H. erato* (n=5) was higher than the 90^th^ quantile of 10,000 permutations, but not in *H. melpomene* (n=2; S.I. Fig S7).

One gene on chromosome 13 stood out, *rugose*, as it was associated with wing shape in both *H. erato* and *H. melpomene*^34^ and overlapped with SHDRs shared in all transects in both species. In *Drosophila* mutants, *rugose* has been shown to affect social interactions, locomotion, and hyperactivity^68^. The highland incipient species of the *H. erato* clade, *H. himera*, has been shown to fly for more hours per day than lowland *H. erato*, suggesting a potentially important role of locomotion to adapt to highland habitats^69^. Additionally, we found that an *H. erato* Eastern SHDR (Fig. 4B) overlapped with a locus recently identified to be differentiated across many pairs of subspecies in several *Heliconius* species and shown to affect wing beat frequency in *Drosophila*^70^. Thus, future studies could focus on functionally testing some of these candidates and ascertain the potentially adaptive functions of candidate regions.

### Same haplotypes underlie parallel adaptation to altitude

High altitude differentiation at the same locus could be driven by the same or different haplotypes under selection. For instance, different *de-novo* mutations at one locus were recently found to confer parallel adaptation to toxic soils in *Arabidopsis*, although most parallel regions were sourced from a common pool of standing alleles^71^. To test whether our candidate regions shared the same haplotypes, we performed local Principal Component Analyses (PCA) with outlier windows of each SHDR (Fig. 5A). While ‘global’ PCAs tend to show relatedness between individuals due to geographic structure or partial reproductive isolation, local PCAs of smaller genomic regions can highlight divergent haplotypes due to, for instance, structural variation or positive selection leading to similar haplotypes in adapted individuals^72,73^. Here we assessed whether genetic variation across individuals at SHDRs (local PCA PC1) could be significantly explained by altitude while accounting for genomewide (‘global’) structuring (Fig. 5 A) to test for evidence for shared allelic basis for altitude adaptation.

**Figure 5.**
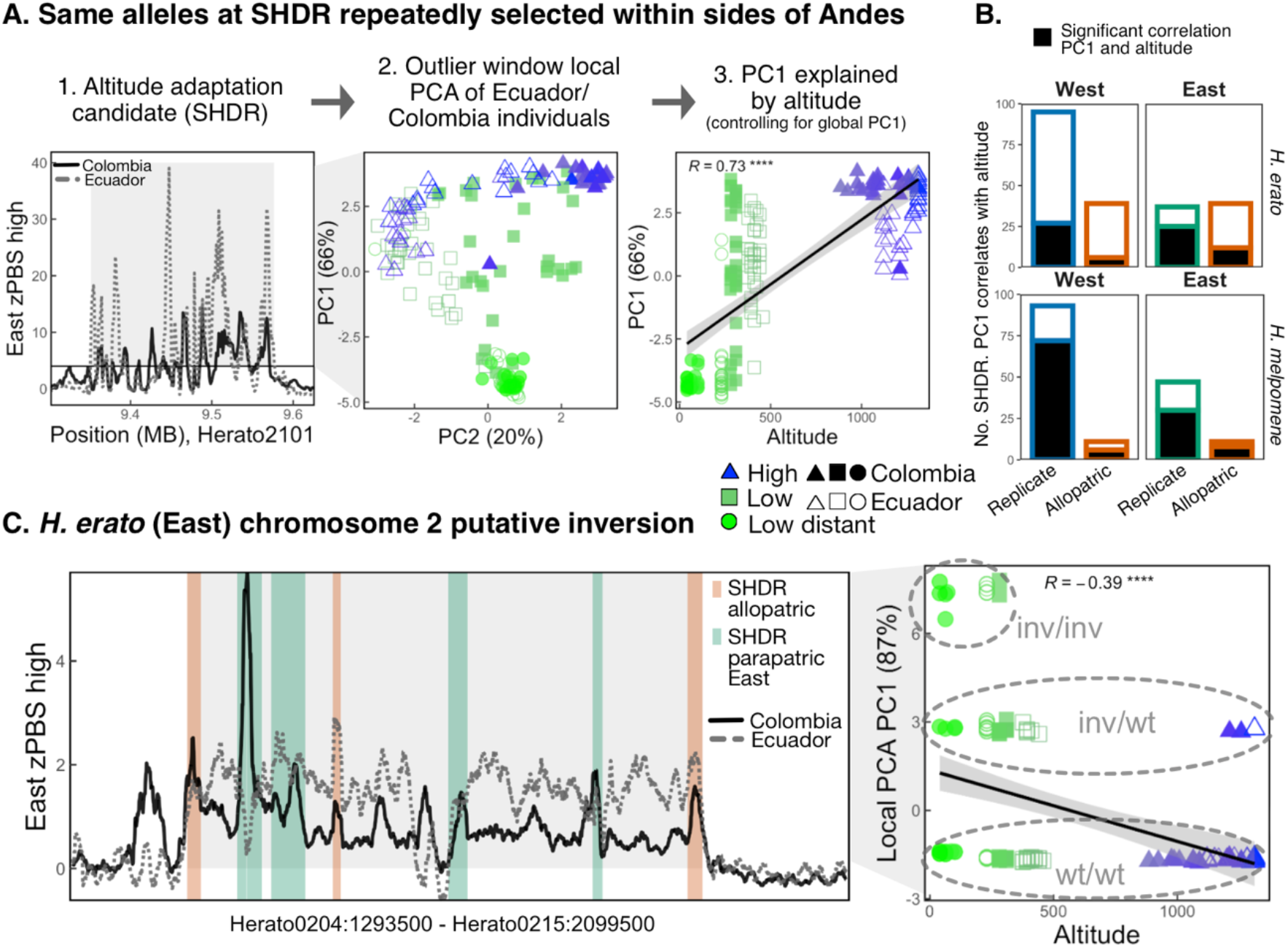
Allele sharing SHDRs and large putative inversion in chromosome 2 of *H. erato* Eastern transects. A) Example analysis to test whether same alleles underlie SHDRs (here depicted *H. erato* Eastern SHDR #77). First, outlier windows (zPBS>4) in either the Eastern Colombia (solid black line) or the Eastern Ecuadorian (dotted grey line) transect are selected (grey panel), and a local PCA with those sites is performed. Then we test whether PC1, the axis explaining most of the variation, is significantly explained by the altitude at which individuals were collected, while controlling for the global PC1 (i.e. neutral population structure). Each point represents an individual, their shape represents transect of origin (Colombia filled, Ecuador empty symbols) and their colour the altitude at which individuals were collected. B) Number of SHDRs where altitude is a significant predictor of local PCA axis 1. C) zPBS highland differentiation in Eastern Colombia (solid black line) and Ecuadorian (dotted grey line) transects across a 6.5 Mbp region of *H. erato* chromosome 2. zPBS lines are regionally smoothed with rolling means of 200 windows, thus some individual outlier windows are higher in value (Fig. 3). SHDRs are shown as vertical segments, colored by whether they represent SHDRs within Eastern transects (green) or allopatric shared across all transects (red, SHDRs shown in Fig. 3). The Pearson correlation coefficient between the putative inversion local PCA PC1 and altitude is shown (****p<0.0001). Homokaryotes for the wildtype arrangement are named wt/wt, heterokaryotes are inv/wt and inversion homokaryotes are labelled inv/inv. The most common arrangement clustered with the Western transects of *H. erato* and outgroups of other species in a neighbour-joining tree, and thus was considered the most likely, non-inverted haplotype (wt/wt) (S.I. Fig. S12).

PCAs in each SHDR were performed with individuals from all elevations in transects connected via gene flow (replicate Colombia/Ecuador transects). Local PCAs at SHDRs often showed individual clustering that differed from the neutral geographic expectations (whole-genome ‘global’ PCAs that included Western or Eastern transects, S.I. Fig. S1), and the first axes of variation tended to explain a much larger proportion of the variation observed (PC1 explained 55% ±20 SD compared to 19% in global PCAs, on average; S.I. Table S3). Out of the four genome-wide PCAs including individuals from replicate transects within sides of the Andes, altitude only explained clustering along PC1 in the Eastern *H. erato* transects (S.I. Table S3, Figure S1). This can be explained by a different highland colour-pattern subspecies in Colombia, *H. e. dignus*, reducing gene flow across the genome (Fig. 2). In contrast, we found that a large proportion of local SHDR PCAs had PC1s that correlated significantly with individual altitude (P<0.05, *H. erato*: East=48%, *H. melpomene*: West=74%, East=66%, S.I. Table S3), except in *H. erato* West where only 25% did (Fig. 5B, S.I. Fig. S8-11). Altitude explained, on average, 12% of the variation in local PC1 while controlling for the global PC1 (altitude partial R^2^, *H. erato*: West= 0.05, East=0.16, *H. melpomene*: West=0.10, East=0.15, S.I. Table S3).

Local PCAs can also highlight divergent haplotypes, putatively associated with inversions, by clustering individuals that possess homozygous or heterozygous haplotypes in those regions^74,75^. We found several *H. erato* SHDRs within a high differentiation block in chromosome 2 (6.5 Mbp, Fig. 5 B), ~0.75 Mbp downstream from a recently identified inversion exclusively present in lowland individuals of southern Ecuador^76^. We performed an additional local PCA across the large putative inversion and found a three-cluster pattern, consistent with the presence of the three inversion genotypes (Fig. 5 B), and a neighbourjoining tree with outgroups supported its appearance in the Eastern lowlands (S.I. Fig. S12). Local PCAs of SHDRs within the inversion region correlated more strongly with altitude than putatively inverted-only haplotypes, indicating that adaptive loci remain differentiated in the highlands and may pre-date the inversion event in the lowlands (S.I. Fig. S8). This is consistent with a model in which the inversion could enhance local adaptation by reducing gene flow between elevations at pre-existing locally-adapted alleles^77^.

Overall, the majority of SHDRs involve the same alleles across transects connected via gene flow. Those SHDRs that do not correlate with altitude could represent cases where different de-novo mutations arose at the same or nearby loci or facilitated by independent introgression events, or indeed may represent regions of differentiation not under positive selection (i.e. false positives). Furthermore, the large putative inversion found exclusively in the lowlands may represent a case of structural variation facilitating adaptation in the highlands^77^. Recent studies on environmental adaptation in seaweed flies and sunflowers, among others, have demonstrated a key role for inversions in maintaining adaptive alleles together and facilitating the evolution of locally adapted ecotypes^72,73^. By studying individual clustering across differentiated loci we have shown that the same alleles often drive parallelisms involved in local adaptation. We thus next turned to identifying the source of the genetic variation causing molecular parallelism across populations.

### The source of parallelism: standing variation and adaptive introgression with high-altitude relatives

The presence of the same putatively adaptive haplotypes on several transects could either reflect: (i) standing variation being repeatedly selected at high altitudes or shared via intraspecific gene flow, or (ii) recruitment of adaptations from other high-altitude adapted species through introgression. With five high-depth individuals per population of *H. erato* and *H. melpomene*, and 116 additional whole-genomes of high-altitude specialist species and outgroups, we tested for signatures of shared standing variation within species and of adaptive introgression in shared high differentiation regions (SHDRs).

To test for excess allele sharing at SHDR, we calculated the F_dM_ statistic in 50 kb windows across the genome^78,79^. For each test, we used a tree with four populations (((P1, P2), P3), O), where P1/P2 reflect the lowland and highland populations, respectively, and P3 is an allopatric high-altitude population or a sympatric high-altitude specialist species (Fig. 6A). Positive F_dM_ values indicate excess allele sharing between P3 and P2 (i.e. between nonsister high-altitude lineages), and negative values indicate excess allele sharing between P3 and P1 (i.e. between non-sister high- and low-altitude lineages, Fig. 6A). We then tested if SHDRs are enriched for outlier positive F_dM_ (i.e. excess allele sharing with the highlands), using the distribution of absolute negative F_dM_ across SHDRs as a null (see Methods for details). This specifically tests whether genomic regions that are differentiated in high-altitude populations (SHDRs) are systematically enriched for alleles shared with allopatric high-altitude populations of the same species or with sympatric specialist high-altitude species.

**Figure 6.**
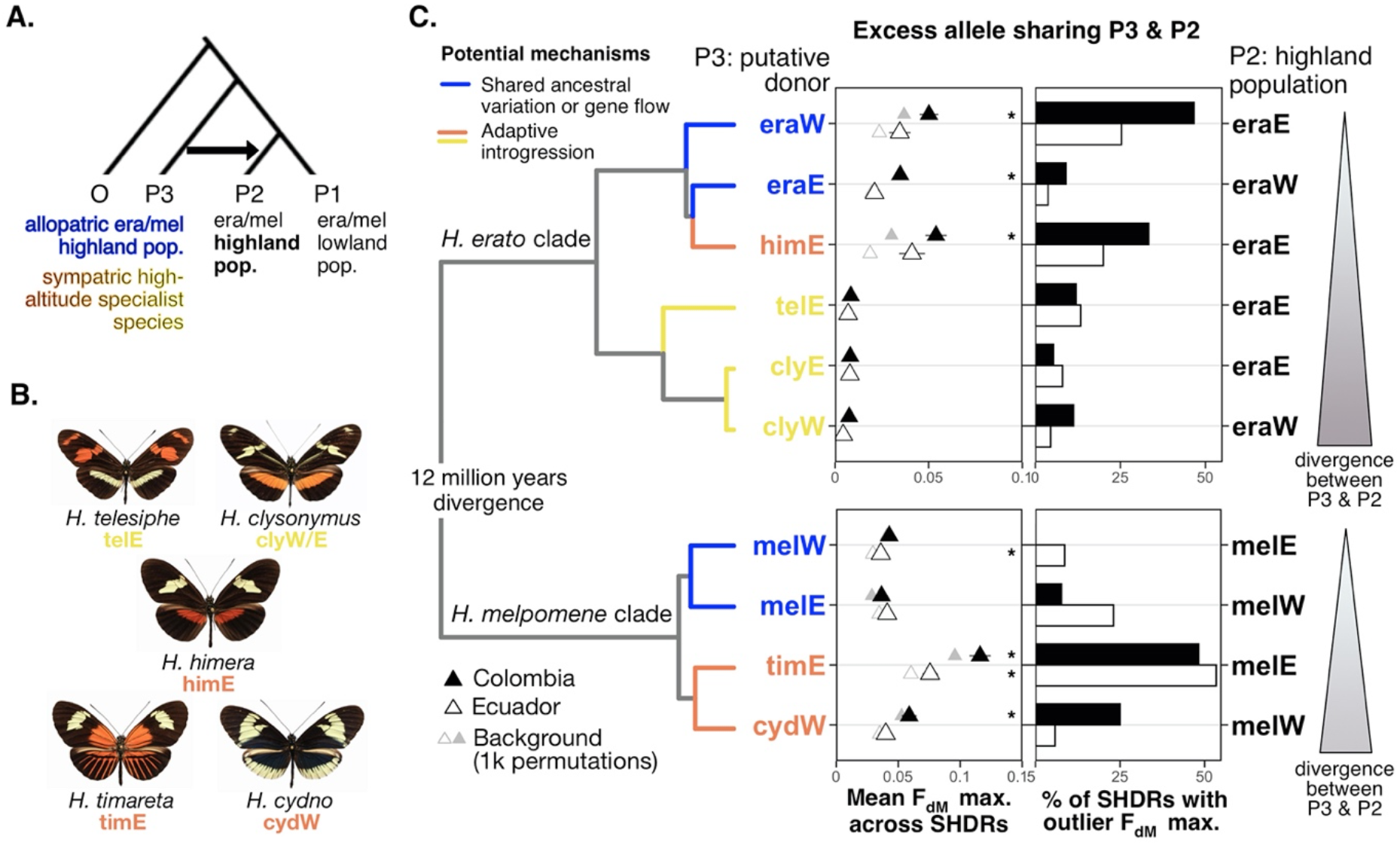
Many SHDRs were sourced from standing variation and putative adaptive introgression from highland-specialist species. A) Tree used to estimate F_dM_ values per 50kb window across the genome in each comparison, which when positive represents excess allele sharing between P2, a highland *H. erato* or *H. melpomene* population, and P3, an allopatric highland population of the same species or a sympatric high-altitude specialist species, compared to a lowland population (P2). Colours of P3 populations or species indicate the potential mechanism driving the excess allele sharing, either intraspecific shared ancestral standing variation (blue) or adaptive introgression from closely (orange) or distantly related (yellow) high-altitude specialist species. B) Excess allele sharing at SHDRs between P3 (putative highland donors, left y axes) and P2 (putative highland recipients, right y axes) across the *H. erato* (top) or *H. melpomene* (bottom) comparisons (phylogeny from Kozak *et al*., 2015)^1^. Left panel shows mean maximum F_dM_ (± S.E.) across SHDRs (western SHDRs if the putative recipient was on the West of the Andes, and vice versa) of the Colombian (solid triangles) and Ecuadorian (unfilled triangles) transects. Background mean maximum F_dM_ values were obtained from 1000 block permutations across the genome and shown in grey. Stars represent comparisons where distribution of maximum F_dM_ (excess allele sharing with the highlands) was significantly higher than absolute minimum F_dM_ (excess allele sharing with the lowlands) distribution across all SHDRs (Kolmogorov-Smirnov tests P<0.05, S.I. Fig. S15-15). Right panels show percentage of SHDRs with evidence of excess allele sharing between P2 and P3, considered significant if they had outlier maximum F_dM_ (>90^th^ percentile of absolute minimum F_dM_ across all SHDRs). Abbreviations not depicted: “era” *H. erato*, “mel” *H. melpomene*. C) Putative donor (P3) high-altitude specialist species.

We first assessed intraspecific allele sharing between allopatric highland populations in transects on opposite sides of the Andes, which split ~0.9 Mya and ~2 Mya in *H. erato* and *H. melpomene*, respectively^36,37^. Signatures of allele sharing likely represent shared ancestral standing variation that pre-dates the Andean split (Fig. 1B), but we cannot rule out gene flow via distant contact zones in the north and south edges of the Andes or periods of secondary contact in the past^37^. Nearly half of *H. erato* Eastern SHDRs had outlier excess allele sharing with the Western highlands in Colombia, whereas only 9% of Western SHDRs did (Fig. 6B, S.I. Fig. S15). Both comparisons resulted in a significant enrichment of excess allele sharing between allopatric highland populations compared to sharing with the lowlands across all SHDRs (Kolmogorov-Smirnov tests P<0.05 as stars in Fig. 6B, S.I. Fig. S15). *H. melpomene* only showed significant enrichment of excess allele sharing between allopatric highland populations of Ecuador in Eastern SHDRs, and the percentage of SHDRs with excess allele sharing was generally lower than in *H. erato* (Fig. 6B). Overall, shared standing variation has been an important mechanism facilitating molecular parallelism in *H. erato*, in which trans-Andean populations share a more recent common ancestor.

We then explored allele sharing between highland populations of *H. erato* and *H. melpomene* and five sympatric high-altitude specialist relatives or those (Fig. 6C). These high altitude species are known to readily or occasionally hybridise with *H. erato* or *H. melpomene*^37,80–82^. To quantify genome-wide evidence of allele sharing we first computed f-branch statistics, which test for gene flow between branches of a phylogeny^83^. As expected, we found evidence of excess allele sharing between all relevant pairs (details in S.I. Note S3, Fig. S13-14)^84^. Excess allele sharing at SHDR between highland populations and sympatric high-altitude specialist species likely represent cases of adaptive introgression. Generally, the more closely related putative donor and recipient species were, the higher proportion of SHDRs that showed excess allele sharing (Fig. 6B, S.I. Fig. S16). For instance, levels of allele sharing in SHDRs were much higher between highland *H. erato* and the closely related *H. himera* than with the distantly related *H. telesiphe* (Fig. 6B). Interestingly, in Eastern *H. melpomene*, SHDR sharing with a closely related sympatric species, *H. timareta*, was more prevalent in SHDR than shared variation with allopatric highland populations of its own species (S.I. Fig. S16D). Admixture between adjacent *H. melpomene* and *H. timareta* populations is well-documented, with strongly selected colour pattern loci having been shared across the species barrier ^81,82,85^

Levels of putative adaptive introgression at SHDRs were also high between *H. himera* and highland *H. erato*, and significantly enriched across SHDRs in Colombia (Fig. 6 B). *H. himera* is a closely related species that split from within the Eastern *H. erato* clade 215,000 - 527,000 years ago^37^, with pre-mating isolation and a range of divergent life-history phenotypes adapted to the highland dry forests it inhabits^86–88^. Admixture is predominantly from *H. himera* into *H. erato*^37^, supporting our hypothesis that the high levels of excess allele sharing at SHDRs between the two may represent cases of adaptive introgression into *H. erato*. Our study is the first to show that potentially adaptive alleles other than colour pattern loci have been shared between high-altitude specialist species and *H. erato* and *H. melpomene*, likely facilitating their expansion into the highlands.

The 6.5Mbp putative inversion detected in chromosome 2 of Eastern *H. erato* individuals showed high levels of allele sharing between highland *H. erato* populations and *H. himera*, whereas there was no excess allele sharing with either the highlands or the lowlands when the putative donor species was a distantly related species (S.I. Fig. S17). A neighbour-joining tree of this region revealed that lowland distant individuals that clustered in the local PCA formed a monophyletic group divergent from all other *H. erato* populations, including allopatric populations in the west of the Andes (S.I. Fig. S12). This suggests that the inversion may have arisen anciently, prior to the western and eastern Andean split of *H. erato*. Its maintenance in the lowland populations may protect locally adaptive alleles from maladaptive migration load and/or promote the accumulation of novel, locally adaptive mutations^73^. Furthermore, its absence in the highlands allows for ongoing gene flow between highland *H. erato* populations and the closely related highland specialist *H. himera*. Supergene evolution in another species of this genus, *H. numata*, has been linked to the introgression of a chromosomal inversion^89^, highlighting the role of structural variation and hybridisation in providing novel genetic architectures that can promote adaptation. Future work could investigate the potential role of this inversion in maintaining locally beneficial allele clusters and their associated adaptive phenotypes.

## Conclusions

By studying recently and anciently diverged populations at different altitudes within and across sides of the Andes of two species we have uncovered (i) strong signatures of high-altitude differentiation in narrow regions across the genome, consistent with positive selection, (ii) high levels of molecular parallelism between transects of the same species but no sharing across species, and (iii) an important role of standing variation and adaptive introgression from high-altitude specialist species in adaptation to these environments. The overall lack of molecular parallelism across species points towards genetic redundancy of polygenic evolution that allows different combinations of alleles to confer adaptation to the same environments^28^. The evolutionary success of *H. erato* and *H. melpomene* in colonising a wide range of elevations has likely been facilitated by abundant genetic diversity, as well as by intra- and interspecific gene flow allowing for the sharing of pre-existing adaptive alleles. Together, our study highlights the value of extensive replication across space and large whole-genome datasets for understanding the molecular underpinnings of local adaptation in the wild. Both standing genetic variation and recent hybridization can supply the selection targets required for adaptation to a new environment, which emphasizes the importance of preserving gene flow and connectivity between populations if organisms are to adapt to everchanging environmental pressures.

## Methods

### STUDY SYSTEM AND WILD BUTTERFLY COLLECTION

*H. erato* and *H. melpomene* can be found across most of the Neotropics and have Müllerian aposematic mimicry to advertise their toxicity to predators, thus share colour pattern when inhabiting the same areas^90^. They can be found continuously coexisting across elevational transects ranging from sea level up to 1600 m along the Andean mountains, and *H. melpomene* can be found across elevations up to 1800 m. Butterflies were collected from 111 different locations, which we grouped into 30 populations, corresponding to four transects Colombia West/East, and Ecuador West/East (Table S1). In each transect populations were either in the highlands (altitude mean=1235 m), lowlands (altitude mean=364 m), or distant lowlands (altitude mean=95 m) to control for genetic drift due to isolation by distance (Fig. 2A). The Andes acts as a barrier to gene flow, as elevations in these latitudes are too high for butterflies to fly across and have been for at least 8 million years^91^, which pre-dates the expansion of both species across these latitudes^36,37^. All but one of these transects (Colombia East) had the same subspecies, i.e. geographic colour morph, in the three elevations, to avoid differentiation due to highly divergent colour pattern loci (Fig. 2A). Detached wings were stored in glassine envelopes and bodies in EtOH (96%) vials. We additionally collected high-altitude specialist relatives of *H. erato* and *H. melpomene* that have potential for admixture between them. The *H. erato* relatives were *H. himera* and *H. telesiphe* from the Eastern Andes, and *H. clysonymus* which is found on both sides of the Andes. The *H. melpomene* relatives were *H. timareta* and *H. cydno*, from the Eastern and Western Andes, respectively. More distantly related outgroups were also sampled, *H. eleuchia* and *H. hecale* for *H. erato* and *H. melpomene*, respectively.

### WHOLE-GENOME SEQUENCING

Whole genome sequence data from 518 individuals was analysed in this study, 444 were newly sequenced here, while the rest were obtained from published studies (n=74). Of the newly sequenced individuals 365 were sequenced at low-medium depth with BGI (~6X), and 79 were sequenced at high depth with Novogene (~18X-30X), at least 5 per population. For the high-altitude specialist species dataset and outgroup species, we obtained high-depth whole genome sequencing data for 116 individuals, 63 of which were newly sequenced for this study at ~20X depth with BGI. A full list of individuals, localitites, and accession numbers can be found in Supplementary Table 1. Individuals with *H. melpomene malleti* phenotypes (Fig. 1) were genotyped with a restriction digest, following Nadeau *et al*. 2014^54^, to identify cryptic individuals of the species *H. timareta ssp. nov*., which are indistinguishable phenotypically from *H. m. malleti*. We extracted DNA with QIAGEN DNeasy Blood and Tissue extraction kits, including RNA removal, and confirmed DNA integrity and concentration (minimum of 10 ng/μL) using Qubit. DNA samples were stored at −20°C until library preparation. For the individuals that were sequenced with low-medium depth, a secondary purification was performed with magnetic SpeedBeads™ (Sigma) and we prepared Nextera DNA libraries (Illumina, Inc.) with purified Tn5 transposase^92^. PCR extension with an i7-index primer (N701–N783) and the N501 i5-index primer was performed to barcode the samples. Library purification and size selection was done using magnetic SpeedBeads™ (Sigma). We confirmed adaptor lengths through TapeStation High sensitivity T1000 (Agilent Technologies, CA, USA) and gel electrophoresis. Pooled libraries were sequenced by The Beijing Genomics Institute (China) using HiSeq X Ten (Illumina). Library preparation and sequencing of the high-depth *H. erato* and *H. melpomene* individuals was carried with HiSeq X platform (150bp paired-end) by Novogene.

### STATISTICAL ANALYSES

All non-genomic analyses were run in R V2.13 (R Development Core Team 2011) and graphics were generated with the package *ggplot2*^93^.

#### Read mapping and genotype calling

We aligned the sequence data of all individuals of the two focal species and their relatives to their corresponding reference genomes, either *H. melpomene* version 2.5^94,95^ or *H. erato demophoon*^55^, obtained from Lepbase^94^, using bwa mem (v 0.7.15 Li, 2013). We used samtools (v 1.9 Li et al., 2009) to sort and index the alignment files. Duplicates were removed using the MarkDuplicates program in Picard tools (v 1.92 Broad Institute, 2018). Genome-wide mean sequencing depth was calculated with samtools (v 1.9 Li et al., 2009). Mean sequencing depth was very similar across areas of *H. erato* (mean=8.93, Fig. S18 and was generally higher for *H. melpomene* (mean= 12.3), but more variable, especially in Colombia where many sequences were obtained from published studies (Supplementary Table 1). Most of the analyses described below for *H. erato* and *H. melpomene* were performed with genotype likelihoods in ANGSD and low or variable sequencing depths are thus accounted for.

However, for our phylogenetic datasets combining our samples with other species for phylogenetic tree reconstruction and tests of admixture, we restricted the *H. erato* and *H. melpomene* samples to the five individuals per population with high sequencing depth. We mapped the high-altitude specialist relatives and outgroups of *H. erato* and *H. melpomene* to the respective reference genomes as explained above (Supplementary Table 1). We used a genotype calling approach with GATK v. 3.7^99^ to obtain a vcf file each for the *H. erato* and *H. melpomene* clade. Genotypes were called with HaplotypeCaller for each individual and variants were then called with GenotypeGVCFs across all individuals combined. The vcf files were filtered with vcftools v. 0.1.15^100^ to remove genotypes with less than 3 reads, monomorphic sites, multi-allelic sites, insertions and deletions (indels), and sites with more than 50% missing data.

#### Isolation by distance and Isolation by environment

To study Isolation by Distance (IBD) and Isolation by Environment (IBE), we first calculated all pairwise genetic differentiation (F_st_) between all populations on each side of the Andes that had at least 5 individuals each, i.e. *H. erato* west (n_population_=7), *H. erato* east (n_population_= 11), *H. melpomene* west (n_population_=7), *H. melpomene* east (n_population_= 9), hereafter “side-species replicate” (Table S1). We calculated pairwise population genetic distance with the function calculate.all.pairwise.F_st_() from the R package BEDASSLE^39^. This requires a matrix of allele count data, with populations as rows and number of biallelic unlinked loci sampled as columns, which we obtained with ANGSD and custom scripts. First, we obtained a list of highly polymorphic SNPs per side-species replicate, by (i) heavily filtering 10 random individuals and obtaining minor allele frequencies (-doMaf 1) by forcing the major allele to match the reference state, so that it is the same across all populations (-doMajorMinor 4) (ii) extracting the sites and the major/minor allele frequencies, (iii) creating an indexed sites file (angsd sites index), (iv) subsetting so that sites are at least 2kb apart, to prune for linkage disequilibrium. This list of sites and regions was then used to obtain minor/major allele frequency counts with all individuals per population and forcing the major/minor allele to match the ones given by the sites file (-doMajorMinor 3). From the resulting allele frequencies per population, we calculated allele counts, by multiplying by the number of individuals per population and the number of chromosomes samples (2, diploid). We obtained the required allele count matrix by concatenating all populations per side-species replicate, and only keeping loci with allele counts for all populations.

Geographical distance between populations measured as a straight line through the landscape is not biologically representative of organisms moving through space. To account for topographic complexity, we obtained topographic least cost paths with the R package *topodistance*^101^. With historical records from the Earthcape database^102^, we created a binary habitat suitability raster based on the elevational range of each species, so that least cost paths between populations never included elevations that these butterflies do not inhabit. Then we used the function topoLCP() to get the least cost path distance between populations (Fig. S3, Wang, 2020), and use this distance as a proxy of isolation by distance between populations.

#### Differentiation and Selection statistics

To search for signatures of local adaptation to high altitude we use a measure of lineage-specific differentiation, population branch statistics (PBS)^44^. PBS is a summary statistic based on pairwise genetic differentiation (Fst) among three populations, two of which are located closely geographically (high, low) and one distant outgroup (low distant). For each population trio (high, low, low distant Fig. 1 A), we computed PBS with ANGSD^103^. We first obtained genotype likelihoods to calculate the site-frequency spectrum (SFS) per population. Then, we computed 2D-SFS for each population pair (hih-low, high-low.distant, low-low.distant) with the function realSFS. We then used the 2D-SFS as a prior for the joint allele frequency probabilities at each site, which are used to compute per-site pairwise Hudson’s Fst^104^ as interpreted by Bhatia (Bhatia et al., 2013, realSFS fst index -whichFst 1) among the three populations (Fst_high-low_, Fst_high-low_, Fst_high-low.distant_) and PBS per population (PBS_high_, PBS_low_, PBS_low.distant_). This is achieved by first transforming pairwise F_st_ values into relative divergence times:

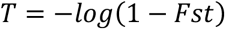

To then obtain PBS for a given population (here the highlands):

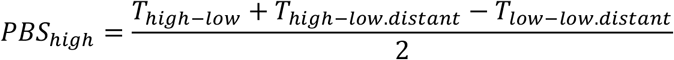

This quantifies the magnitude of allele frequency change in the highland lineage since its divergence from the lowland and lowland distant populations. Regions of the genome with large PBS values represent loci that have undergone population-specific sequence differentiation consistent with positive selection. We computed weighted Fst and PBS averages per 5kb window size and 1kb steps (realSFS fst stats2), with the same window centres across datasets (-type 0). For transects where only two out of three populations had any individuals (Colombia west for *H. erato* and Ecuador west for *H. melpomene* Fig. 1 A), we computed pairwise Fst only. Finally, we normalized PBS and Fst window values with z-scores, i.e. number of standard deviations from the mean, so that divergence is comparable across transects.

#### HDR parallelism between replicates

To measure genetic parallelisms in adaptation to high altitude, we quantified overlap of outlier windows and adjacent regions, i.e. high differentiation regions (HDR), across replicate and allopatric transects and species. We considered outlier windows to be those with values above 4 standard deviations from their mean (zPBS_high_>4, zFst>4). We then expanded outlier window positions 50kb upstream and 50kb downstream from the window center (HDR) and checked for overlaps with other transects, either within or across sides of the Andes for each species, to highlight parallelism when visualizing patterns of genome-wide divergence. High-differentiation regions with any overlap with other transects are termed shared HDRs (SHDRs). To check for HDR overlaps between species, we mapped the *H. melpomene* windows (starts and end positions) to the *H. erato* reference genome using a chainfile from Meier et al. (2021)^76^ and the liftover utility (Hinrichs, 2006).

To test whether the level of parallelism observed between transects within and across sides of the Andes was higher than expected by chance, we used the R package intervals^106^. We first created outlier window intervals (HDRs), by obtaining the start and end positions of continuous blocks of outlier windows (±50kb buffers) with the function Intervals() (options type=”Z”, closed). We obtained the observed proportion of total intervals that overlapped, at any of their positions, with outlier intervals in the other transect within sides of the Andes, or with outlier intervals in both, the other transect within the same side of the Andes and the two transects on the other side of the Andes (allopatric sharing). We calculated confidence intervals around the proportion of overlapping intervals by performing random jacknife block resampling across the genome. We then simulated 10,000 randomized distributions of outlier-window intervals across the genome per transect, per species (n=8). In each simulation set, we randomly placed the same number of HDR intervals and of the same size as the observed outlier-window intervals for those transects. With these, we estimated the proportion of simulated intervals that overlapped with observed HDR outlier intervals within and across sides of the Andes, obtaining as a result a null distribution of interval overlap proportions.

#### Measures of nucleotide diversity, selective sweeps, and recombination rate

We studied genetic variation within and across populations by deriving three summary statistics from ANGSD thetas estimations^103^, Tajima’s D, which estimates the deviation of a sequence from neutrality, nucleotide diversity (pi, or population mutation rate), and D_xy_ or absolute divergence, which calculates pairwise differences between sequences of two populations excluding differences between sequences within populations. ANGSD has been found to be an accurate estimator of nucleotide diversity because it includes invariant sites^107^. Firstly, we obtained folded global site-frequency spectra for each population. Then we calculated pairwise nucleotide diversity per site (thetaD, realSFS saf2theta). Finally, we performed sliding window analysis of 5kb window size and 1kb steps (thetaStat do_stat) to obtain sum of pairwise differences, Tajima’s D, and total effective number of sites per window. Nucleotide diversity (pi) was obtained by dividing the sum of pairwise differences by the total number of sites per window. For all transects, we calculated the difference in nucleotide diversity per window between highland and lowland distant populations (or low populations if low distant individuals unavailable), which is expected to be negative if a selective sweep led to locally reduced diversity in highland populations. Absolute divergence (D_xy_, Nei, 1978) between high (population A) and low/low distant (population B) populations was estimated by additionally obtaining pairwise nucleotide diversity per site (thetaD) for all individuals pooled from populations A and B (population AB), and then per-site D_xy_ obtained:

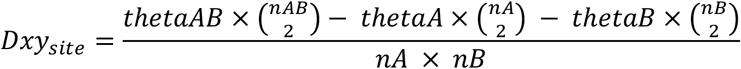

With n being the number of individuals per population (A, B, or pooled AB) and thetaD obtained from realSFS saf2theta. Mean D_xy_ was estimated for the same 5kb windows with 1kb steps. Finally, we obtained recombination rate for 50kb windows along the genome from the mean population recombination rate (*ρ* = 4*N_e_r*; *r* = probability of recombination per generation per bp) estimated for 13 *H. erato* populations from across the range in a recent study^109^ and for 100kb windows of four *H. melpomene* populations^67^. Note that the *H. erato* genome is larger than *H. melpomene* (383 Mb and 275 Mb, respectively), hence the difference in window sizes.

#### Testing for significance of selection statistics in SHDRs

To test for positive selection we assessed Tajima’s D, difference in nucleotide diversity across elevations Δπ (π_high_-π_low_), and absolute divergence (D_xy_) within SHDRs and compared values to simulated distributions. We used the same permutation approach described for assessing HDR parallelism, by randomly placing the same number of intervals and of the same size as the observed HDRs for each transect 10000 times. We then obtained minimum Tajima’s D and Δπ and maximum D_xy_ within each simulated SHDR and permutation, and only considered a SHDR to be an outlier for a given selection statistic if the observed maximum (Tajima’s D, Δπ) or minimum (D_xy_) value was above the 90^th^ or below the 10^th^ percentile of simulated values. Number of outlier selection statistics per SHDR was tallied and compared across replicate or allopatric sharing in each species.

#### Global and local PCAs

To assess neutral genetic variation between individuals and populations, we performed principal component analysis (PCA) in the eastern and western transects separately for each species, i.e. two transects per PCA (Fig. 1 A). We first obtained a random subsample of 10% of windows that did not have high differentiation across populations, i.e. with zPBS/zFst < 4, and then pruned for linked sites by only retaining 1 site for every 10kb, yielding 14995 and 8293 sites for *H. erato* and *H. melpomene*, respectively. We used the program ANGSD (v 0.933, Korneliussen et al., 2014) to obtain genotype likelihoods in beagle format (-doGlf 2) for all individuals. In *H. erato*, we excluded chromosome 2 as it contains a large inversion which could distort the neutral differentiation landscape. Genotype likelihoods were used as input for PCAngsd, which incorporates genotype uncertainty from genotype likelihoods to obtain a covariance matrix across all individuals^110^.

To assess whether the same haplotypes were involved in adaptation to altitude across replicate transects (within sides of the Andes), we perfomed local PCAs with outlier windows (zPBS/zFst > 4, i.e. >4 standard deviations from the mean) of each SHDR (total = 370 local PCAs). All individuals from replicate transects were included, leading to local PCAs for all Western and Eastern SHDRs of each species that included Colombia and Ecuador samples. We obtained genotype likelihoods in beagle format (-doGlf 2) as input for PCAngsd, similarly to the population structure analysis. We then assessed whether altitude was a significant predictor of individual clustering in each SHDR by building a linear model where local PCA PC1 was the response variable, and altitude and global (genome wide) PCA PC1 the predictors, to account for population structure. We considered that individuals in each replicate transect had the same haplotypes in SHDRs if altitude was a significant predictor of local PCA PC1, while controlling for global PC1 (Fig. 5). We additionally obtained the overall variation explained by the fitted linear models (R^2^) for each SHDR local PCA and the relative contributions of each explanatory variable (altitude and global PCA PC1, partial R^2^), estimated with the package relaimpo^111^.

#### Measures of excess allele sharing

We used *ABBA-BABA*-related statistics to examine patterns of allele sharing between closely or distantly related high-altitude species and our study *H. erato* and *H. melpomene* populations. These statistics test for an excess of shared derived variation between lineages to distinguish gene flow or ancestral population structure from the incomplete lineage sorting (ILS) that can occur during a simple tree-like branching process. To examine genome-wide patterns of excess allele sharing between populations and species, we obtained F branch statistics implemented with the package *Dsuite*^83^. Fbranch summarises and visualises patterns of excess allele sharing across phylogenetic datasets. We performed linkage-pruning to obtain a genome-wide average of excess allele sharing. Using a custom script (https://github.com/joanam/scripts/blob/master/ldPruning.sh) we removed sites above an LD-threshold of R^2^>0.1 with plink v. 1.07^112^. To reconstruct the backbone phylogeny for excess allele sharing tests, we extracted for each population or species the individual with highest sequencing depth from the vcf file using vcftools v. 0.1.15^100^. The vcf file was then converted to phylip with a custom script (https://github.com/joanam/scripts/blob/master/vcf2phylip.py). We reconstructed the phylogeny of the melpomene and erato clade separately with RAxML v. 8.2.9 using the GTRGAMMA model. Using this backbone tree, we used the LD-pruned vcf files of all melpomene/erato clade individuals to computed f statistics (tests of excess allele sharing) across all possible sets of three populations or species with Dsuite Dtrios. Next, we summarized these statistics with Dsuite Fbranch. The extent of gene flow in the eastern Andes between *H. melpomene* and *H. timareta* can skew genome-wide trees. Thus, to assess the levels of gene flow, we constrained the *H. melpomene* clade to be monophyletic following the species tree. In order to remove spurious signatures of excess allele sharing that are not significant, we set Fbranch values to 0 if the z-score was greater than 3 with a custom script (https://github.com/joanam/scripts/blob/master/removeNonsignDsuite.r). Lastly, we plotted the Fbranch statistics along the phylogeny with dtools.py of the Dsuite package.

To test for adaptive introgression from high-altitude specialist species into *H. erato* and *H. melpomene* highland populations, we computed *f*_dM_, a statistic of excess allele sharing suitable for small genomic regions. This test is based on a set of four populations, including two sister taxa (P1 & P2), a close relative (P3) that may have admixed with one of these sister taxa and an outgroup (O). Here, the sister taxa represent the *H. erato* or *H. melpomene* lowland (P1) and highland populations (P2), whereas P3 represents a highland specialist species that may have contributed beneficial gene variants to the *H. erato* and *H. melpomene* highland populations. f_dM_ quantifies gene flow between P3 and P2 or between P3 and P1. In addition, we ran these tests with the allopatric erato/melpomene populations as P3, to test if at SHDRs, the same haplotypes are found in high-altitude populations on both sides of the Andes, potentially due to parallel selection on the same haplotypes. We estimated f_dM_ for non-overlapping 50 kb windows across the genome with the ABBABABAwindows.py script by (Martin et al., 2014) from https://github.com/simonhmartin/genomics_general. We considered individual SHDRs as f_dM_ outliers if their observed maximum f_dM_ value was >90^th^ percentile of the absolute minimum f_dM_ values across all SHDRs. Additionally, we tested for overall enrichment of excess allele sharing between P3 and P2 (i.e. with the highlands) across all SHDRs, by testing with a Kolmogorov-Smirnov tests if the distribution of maximum f_dM_ values across all SHDRs (i.e. allele sharing with the highlands, P2) was significantly higher than the absolute minimum f_dM_ values across all SHDRs (i.e. allele sharing with the lowlands, P1). This was repeated for each Colombia/Ecuadorian clines with their respective potential donors (P3).

## Acknowledgements

We are grateful to all the field assistants who have collected samples for this study, Narupa Reserve (Jocotoco Foundation, Ecuador), Jatun Satcha Reserve (Ecuador), and Universidad Regional Amazónica Ikiam for their support. We thank Steven van Belleghem for sharing recombination rates data for *H. erato*, and Emma Curran and Juan Enciso for providing some of the genomic sequences, and the Butterfly Genetics Lab (Cambridge) for helpful feedback.

G.M.-K. was supported by a Natural Environment Research Council Doctoral Training Partnership (NE/L002507/1). This work, N.J.N. and C.D.J. were supported by the Natural Environment Research Council (grant number: NE/R010331/1) and by a European Research Council Grant (339873) to C.D.J.. Some of the sequence data was generated under a NERC fellowship (NE/K008498/1) to NJN. Funding was provided to C.N.B. by the Spanish Agency for International Development Cooperation (AECID, grant number 2018SPE0000400194). C.S and NR were funded by Fondos Concursables Big - grant IV-FGD005/IV-FGI006 Universidad del Rosario. SHM was supported by a NERC IRF (NE/N014936/1). Y.F.C. was supported by the European Research Council Starting Grant 639096 “HybridMiX” and the Max Planck Society. Open access funding provided by University of Cambridge. Deposited in PMC for immediate release.

In Colombia, field collections were conducted under permit no. 530 issued by the Autoridad Nacional de 539 Licencias Ambientales of Colombia (ANLA). In Ecuador, collections during November-December 2011, and September-October 2012, were done under permit 0033-FAU-MAE-DPO-PNY and exported under permits 001-FAU-MAE-DPO-PNY and 006-EXP-CIEN-FAU-DPO-PNY. Permits were obtained from Parque Nacional Yasuní, Ministerio Del Ambiente, La Dirección Provincial de Orellana. Collections in Ecuador during 2017-2019 were conducted under the permit provided by the Ministerio del Ambiente, Ecuador (MAE-DNB-CM-2017-0058).

## Data availability

Sequence data is deposited at the NCBI Short Read Archive (primary accession: PRJEB36288) or elsewhere on ENA, as specified in Supplementary Table 1. Scripts have been made available in the public repository Zenodo (TBC). All records associated to the individuals used for this study are available in the *Heliconius* Earthcape database (https://heliconius.ecdb.io/, Jiggins et al., 2019).

