## Supplementary Information for "Repeated genetic adaptation to high altitude in two tropical butterflies"

### Note S1. HDR summaries and testing impact of buffer size around outlier windows on parallelism

In the three *H. erato* clines with three populations (PBS), we found an average of 216 HDRs (±47) covering 6.4% of the genome (±1.2%), including upstream and downstream region buffers of 50kb. In the three *H. melpomene* clines with PBS, we found 242 HDRs (±65) covering 10.9% of the genome (±2.5%), on average. The clines for which only two populations were sampled (Fst), had a higher number of HDRs, 400 and 405 HDRs, covering 11.4% and 17.1% of the genome for *H. erato* and *H. melpomene*, respectively.

We tested the effect of changing buffer size on the difference between the observed and simulated proportion of HDRs shared across clines. We found that this difference was generally robust to the size of the buffer, except when increasing the buffer to ±100kb in *H. melpomene*, which led to a larger difference between the observed and simulated values (S.I., Fig. S18). Thus, we used ±50kb as buffers around outlier windows, which is conservative given that well-studied regulatory regions can be up to 100kb away of positively selected colour pattern loci in these species[1].

**Note S2. East SHDRs test for additional signatures of selection in published high-quality linked-read dataset**

To further validate signatures of selective sweeps in our altitude Shared High Differentiation Regions (SHDRs), we used a recently published linked-read dataset to run a Genome-wide association study with individual altitude as a phenotype and tested for enrichment of selection statistics within SHDRs (Fig. S5). Four selection statistics were calculated for the highland and lowland populations in Meier *et al*. 2021[2], integrated haplotype score (iHS), nucleotide diversity ($\Pi$) across elevations ($\Delta\Pi = \Pi_{high} - \Pi_{low}$), omega ($\omega$), and composite likelihood ratio to detect selective sweeps using sweeD. We additionally performed a genome-wide association study with the same dataset, where the phenotype/response variable was set as the altitude at which individuals had been collected while controlling for sex, wing area, and admixture proportions as covariates. We followed the methods and outlier association region detection pipelines outlined in Montejo-Kovacevich et al. 2021 [3], which used this sequencing dataset to investigate wing shape genomics.

We found that over half of eastern SHDRs of both *H. erato* and melpomene overlapped with at least one selection outlier or GWAS association region of this independent altitudinal re-sequenced cline (Fig. S5 B, C). As expected, the number of western SHDRs that overlapped with selection outliers was lower, as the dataset from Meier *et al.* 2021 is from individuals collected on a cline on the East of the Andes (Fig. S5 A). We found that 50% and 15% of eastern parapatric SHDR in *H. erato* and melpomene, respectively, overlapped with strongly altitude-associated regions in the haplotagging GWAS, which make up 0.44% of the genome, whereas only 8% and 11% of parapatric western SHDRs overlapped with GWAS outliers (Fig. S5 D, E). Allopatric SHDRs showed high levels of overlap with haplotagging-derived selection statistics in all clines (52% ±15 SD, on average)

**Note S3. Genome-wide evidence of admixture across clades**

We assessed genome-wide evidence of ancestral standing variation or admixture by obtaining f-branch statistics from a balanced dataset of five high depth samples per population or species. In the *H. erato* clade, we found strong evidence of genome-wide allele sharing between the highland *H. telesiphe* / *H. clysonymus* clade and the western *H. erato* clade (mean fbranch= 0.034, Fig. S13). There was excess allele sharing between geographically isolated western and eastern highland populations of *H. erato*, especially in the Colombia western populations (Fig. S13), indicating the presence of ancestral standing variation as gene flow is absent. Furthermore, the highland species *H. clysonymus*, which is found on both sides of the Andes, had excess allele sharing with all populations of the *H. erato* clade, including the incipient eastern species *H. himera* (mean fbranch= 0.043, Fig. S13, Fig. 6). In *H. melpomene*, fbranch values were generally higher, probably because of a more closely related set of species and populations, leading to higher admixture (Fig. S14). Contrary to expectations, in H. melpomene the strongest signatures of genome-wide excess allele sharing corresponded to comparisons with sympatric high-altitude closely related species rather than within-species comparisons between eastern and western clades of the Andes (Fig. S14). These results support the presence of allele sharing between high-altitude specialist species and the *H. erato* and *H. melpomene* populations under study, which could allow for locally adaptive genomic regions to have been shared via adaptive introgression.

**Table S1.** Population summaries (n=30) corresponding to the points shown in Fig 1A. Nearby localities were considered one population, individual localities listed as they appear on the Earthcape database separated by ";" [4].

| Name | Side Andes | Country | Alt. | Localities on Earthcape | # *H. erato* | # H. melp. | latitude | longitude | altitude |
|---|---|---|---|---|---|---|---|---|---|
| flo | East | Colombia | High | Sucre, Florencia; Sucre; Doraditas; Finca C. Piñacué; Quebrada Las Doraditas | 4 | 8 | 1.795 | -75.660 | 1174 |
| moc | East | Colombia | High | Mocoa-Subundoy site 4; Mocoa-Subundoy site 5 | 6 | 0 | 1.079 | -76.716 | 1235 |
| cam | East | Colombia | High | Campo Cana 02; Campo Cana 01; Campo Cana 06; Campo Cana 03; Campo Cana 04; Campo Cana 07; Campo Cana 09; Campo Cana Mauricio3; Campo Cana 13; Campo Cana 12; Campo Cana 10; Campo Cana; Campo Cana Mauricio1; Campo Cana 11; Campo Cana Mauricio2 | 14 | 17 | 1.214 | -76.685 | 993 |
| bae | East | Ecuador | High | Km 119 Baeza - Lago Agrio; Baeza - Lago Agrio, Rio Salado | 0 | 10 | -0.191 | -77.697 | 1360 |
| sum | East | Ecuador | High | Wild Sumaco Hummingbird Trail | 0 | 5 | -0.677 | -77.599 | 1468 |
| rev | East | Ecuador | High | Reventador road; Reventador-Sucumbios; San Raphael Falls, | 14 | 3 | -0.030 | -77.529 | 1312 |
| nar | East | Ecuador | High | Baeza-Lago Agrio, Reserva Narupa, P10; Reserva Narupa, bridge; Jondachi, Napo, Ecuador; Mina Negra; Finca Narupa, bridge; Finca Narupa; Finca Jose Simbanas P13; Road after y in km24 Tena-Quito | 18 | 7 | -0.703 | -77.753 | 1178 |
| bra | West | Colombia | High | Rio Bravo - Calima; Rio Bravo 3; Rio Bravo-Calima; Rio Calima 5km | 0 | 16 | 3.632 | -76.713 | 1334 |
| que | West | Colombia | High | Queremal; Queremal (Rio San Juan, Vereda El Digua) | 21 | 2 | 3.504 | -76.757 | 1192 |
| min | West | Ecuador | High | Casa Amarilla, Mindo; Mindo to Finca Birdadvenrure 4; Mindo to Finca Birdadvenrure 1; Mindo to Finca Birdadvenrure 2 | 11 | 1 | -0.055 | -78.785 | 1245 |
| pac | West | Ecuador | High | Nanegal to Marianita; Pacto- Rio Toali, site 4; Pacto to Paraiso, Rio3; Pacto to Paraiso, Rio1; Pacto- Rio Toali, site PAC4; Pacto to Paraiso, Rio2; Pacto- Rio Toali, site 5; Pacto- Rio Toali, site 2; Pacto- Rio Toali, site 6; Pacto- Rio Toali, site 1; Pacto to Paraiso 1 ; Road to Mashpi (116); Mashpi to Pachijal 2; Road to Mashpi 3 (116); Road to Mashpi 2 (116) | 20 | 28 | 0.154 | -78.760 | 1097 |
| fll | East | Colombia | Low | Quebrada La Yuca | 0 | 1 | 1.610 | -75.667 | 340 |

| | | | | | | | | | |
|---|---|---|---|---|---|---|---|---|---|
| guz | East | Colombia | Low | Pto Guzman; finca la cimita Guzman | 12 | 7 | 0.956 | -76.409 | 281 |
| esc | East | Colombia | Low | Finca el Escondite; Via a la Joya-finca el Escondite-vereda san Rafael; Via a la Joya-finca el Escondite-vereda san Rafael | 21 | 2 | 0.795 | -76.585 | 304 |
| jat | East | Ecuador | Low | Limoncocha - El Carmen 2; Limoncocha - El Carmen 1; San Pedro de Arajuno, Río Arajuno; San Pedro de Arajuno, trail; Jatun Satcha P03; Jatun Satcha, jardin botanico; Puni Bocana, trail; Puni Bocana, lodge; Y de Misahuallí; Jardin Aleman, Misahualli; Pununo; Road to Shalcana 2; Road to Shalcana | 33 | 25 | -1.078 | -77.624 | 414 |
| bar | West | Colombia | Low | Santa Barbara | 0 | 1 | 3.830 | -76.786 | 282 |
| cau | West | Colombia | Low | La Elsa - Rio Digua | 0 | 7 | 3.580 | -76.862 | 346 |
| all | West | Ecuador | Low | Alluriquin | 0 | 2 | -0.320 | -79.337 | 300 |
| gua | West | Ecuador | Low | Guayllabamba | 0 | 12 | 0.190 | -78.902 | 532 |
| tor | West | Ecuador | Low | Mashpi town (117); El Tortugo, P28; El Tortugo, P26; El Tortugo, P25; El Tortugo, P24; Pachijal to Tortugo 6 | 32 | 11 | 0.204 | -78.943 | 480 |
| amz | East | Colombia | Low distant | Puerto Nari_o-Quebrada aguas rojas; Puerto Nari–o- Quebrada aguas rojas | 6 | 1 | -3.770 | -70.340 | 101 |
| let | East | Colombia | Low distant | Leticia - Vereda San Jose; Leticia - Km 9.3; Leticia - Reserva Cerca viva km12; Leticia - Tio Tacana 2km | 16 | 7 | -4.147 | -69.960 | 50 |
| ana | East | Ecuador | Low distant | Anangu Boca del Rio ECD OR; Napo Wildlife Center community Anangu | 14 | 5 | -0.504 | -76.388 | 228 |
| yas | East | Ecuador | Low distant | Yasuni Research Station, | 14 | 0 | -0.674 | -76.397 | 230 |
| pot | West | Colombia | Low distant | Potes 2 ; Playa Flores 3; Playa Flores; Cocalito; Cocalito 2 | 0 | 13 | 6.375 | -77.387 | 30 |
| ama | West | Colombia | Low distant | Amargal Station; Amargal 1 | 4 | 5 | 5.572 | -77.502 | 72 |
| ped | West | Colombia | Low distant | San Pedro 1 | 4 | 0 | 3.837 | -77.257 | 4 |
| lad | West | Colombia | Low distant | Ladrilleros; La Barra | 16 | 4 | 3.958 | -77.373 | 34 |
| pue | West | Ecuador | Low distant | Puerto Quito, La Isla | 16 | 0 | 0.110 | -79.230 | 130 |
| ton | West | Ecuador | Low distant | Tonsupa 1 | 22 | 0 | 0.854 | -79.804 | 77 |

**Table S2.** Mean genome-wide Tajima's and nucleotide diversity (pi) per population, calculated in 5kb windows with 1kb steps.

| Species | Pop. | pi | Tajima's D | Cline | Altitude |
|---|---|---|---|---|---|
| *H. erato* | que | 0.024 | -0.444 | Col. West | High |
| | min | 0.022 | -0.504 | Ec. West | High |
| | pac | 0.023 | -0.670 | Ec. West | High |
| | tor | 0.023 | -0.834 | Ec. West | Low |
| | lad | 0.023 | -0.775 | Col. West | Low distant |
| | pue | 0.023 | -0.708 | Ec. West | Low distant |
| | ton | 0.023 | -0.734 | Ec. West | Low distant |
| | cam | 0.032 | -0.751 | Col. East | High |
| | moc | 0.031 | -0.580 | Col. East | High |
| | nar | 0.032 | -1.135 | Ec. East | High |
| | rev | 0.032 | -1.015 | Ec. East | High |
| | esc | 0.033 | -1.317 | Col. East | Low |
| | guz | 0.033 | -1.146 | Col. East | Low |
| | jat | 0.033 | -1.392 | Ec. East | Low |
| | amz | 0.032 | -0.851 | Col. East | Low distant |
| | let | 0.034 | -1.264 | Col. East | Low distant |
| | ana | 0.033 | -1.267 | Ec. East | Low distant |
| | yas | 0.034 | -1.210 | Ec. East | Low distant |
| *H. melpomene* | pac | 0.020 | -0.065 | Ec. West | High |
| | bra | 0.019 | 0.301 | Col. West | High |
| | gua | 0.020 | 0.067 | Ec. West | Low |
| | tor | 0.019 | -0.068 | Ec. West | Low |
| | cau | 0.020 | 0.170 | Col. West | Low |
| | ama | 0.020 | 0.068 | Col. West | Low distant |
| | pot | 0.021 | -0.221 | Col. West | Low distant |
| | bae | 0.019 | -0.260 | Ec. East | High |
| | nar | 0.021 | -0.566 | Ec. East | High |
| | sum | 0.020 | -0.278 | Ec. East | High |
| | cam | 0.021 | -0.610 | Col. East | High |
| | flo | 0.021 | -0.391 | Col. East | High |
| | jat | 0.021 | -1.061 | Ec. East | Low |
| | guz | 0.020 | -0.479 | Col. East | Low |
| | ana | 0.021 | -0.522 | Ec. East | Low distant |
| | let | 0.021 | -0.561 | Col. East | Low distant |

**Table S3.** Local PCA summaries across different types of SHDRs. Linear models were built with local PCA PC1 as a response variable and individual altitude and global (genome-wide) neutral PCA PC1 as explanatory variables, i.e. accounting for genome-wide population structure. Partial $R^2$ for altitude was calculated with the package *relaimpo* [5].

| SHDR type | No. SHDR (no. that are also allopatric SHDRs) | % SHDR with altitude as a significant predictor of local PCA PC1 | % allopatric SHDR with significant correlation between PC1 and altitude | Mean % of variation explained by local PC1 (global PC1, for comparison) | Mean altitude partial $R^2$ for models where altitude was a significant predictor of local PCA PC1 (altitude partial $R^2$ in global PC1 model, for comparison) |
|---|---|---|---|---|---|
| *H. erato* West | 133 (38) | 25% | 16% | 58% (21%) | 0.05 (n.s.) |
| *H. erato* East | 75 (38) | 48% | 32% | 55% (8.8%) | 0.16 (0.11) |
| H. melpomene West | 104 (11) | 74% | 55% | 56% (36%) | 0.10 (n.s.) |
| H. melpomene East | 58 (11) | 66% | 73% | 66% (11%) | 0.15 (n.s.) |

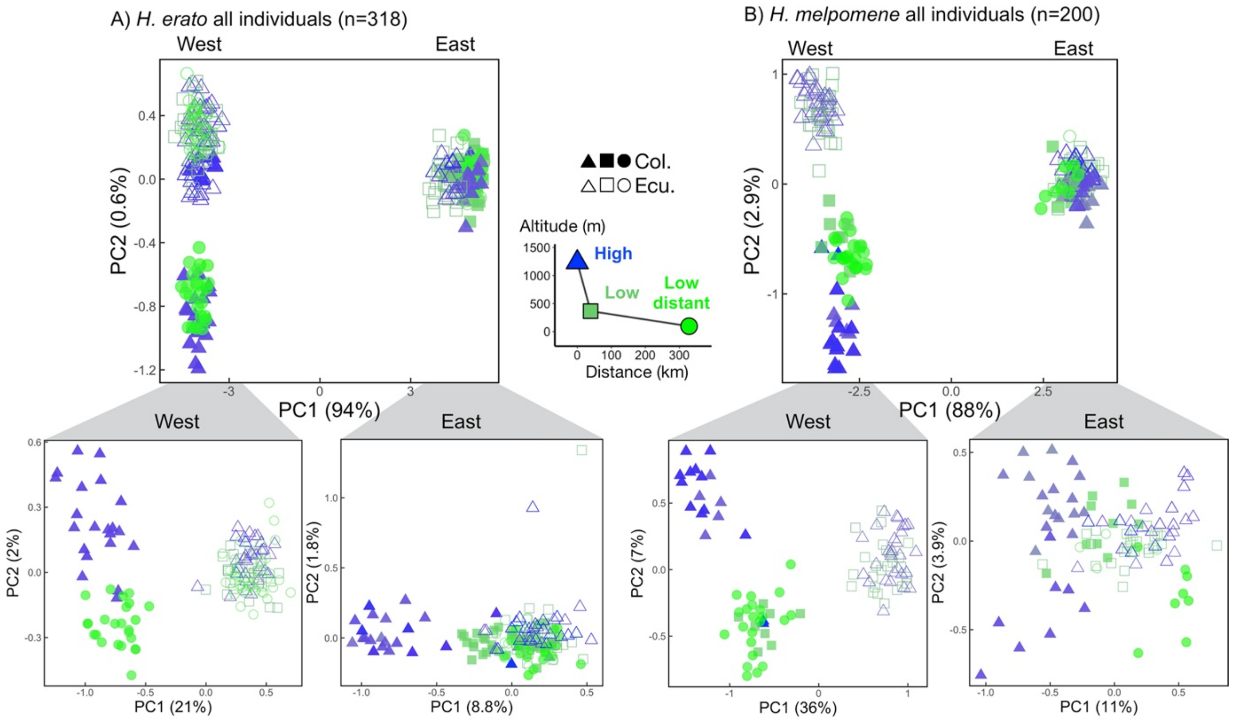


**Figure S1.** Population structure between all individuals of *H. erato* and *H. melpomene* (top panel) and divided into sides of the Andes (bottom panel). Genetic variation along the first two eigenvectors of a principal component analysis (PCA) and the percentage of variance explained by the principal components is shown in brackets. Note that in the West, Ecuador and Colombia have different subspecies/colour patterns (Fig. 1D), and, additionally, the highlands of Colombia East have (filled blue triangles) different subspecies compared to the lowlands.

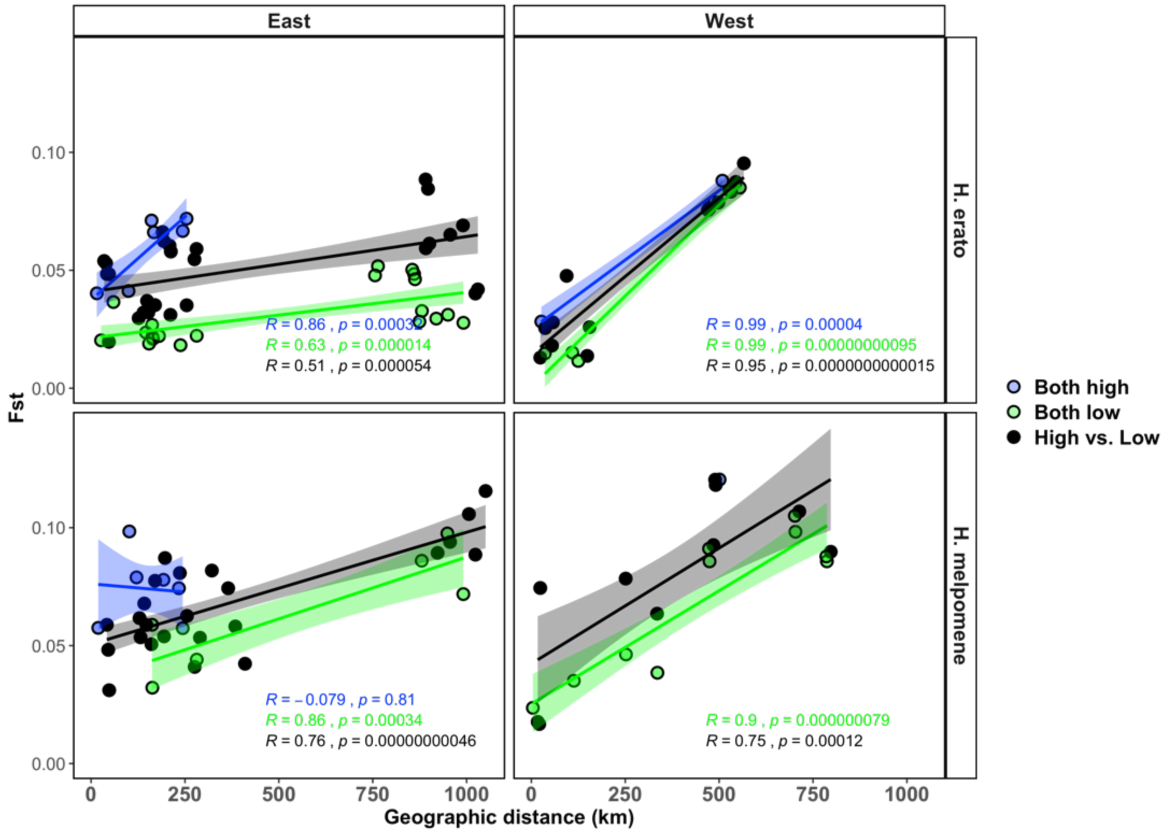


**Figure S2.** Pairwise population genetic differentiation (Fst) between all population pairs per side of the Andes and per species. Population pairs can be both at high altitude (blue), both at low altitude (green), and high vs. low altitude (black). Geographic distance incorporates topography and least cost paths to circumvent elevations at which these species are not found. Pearson correlation and p-values shown.

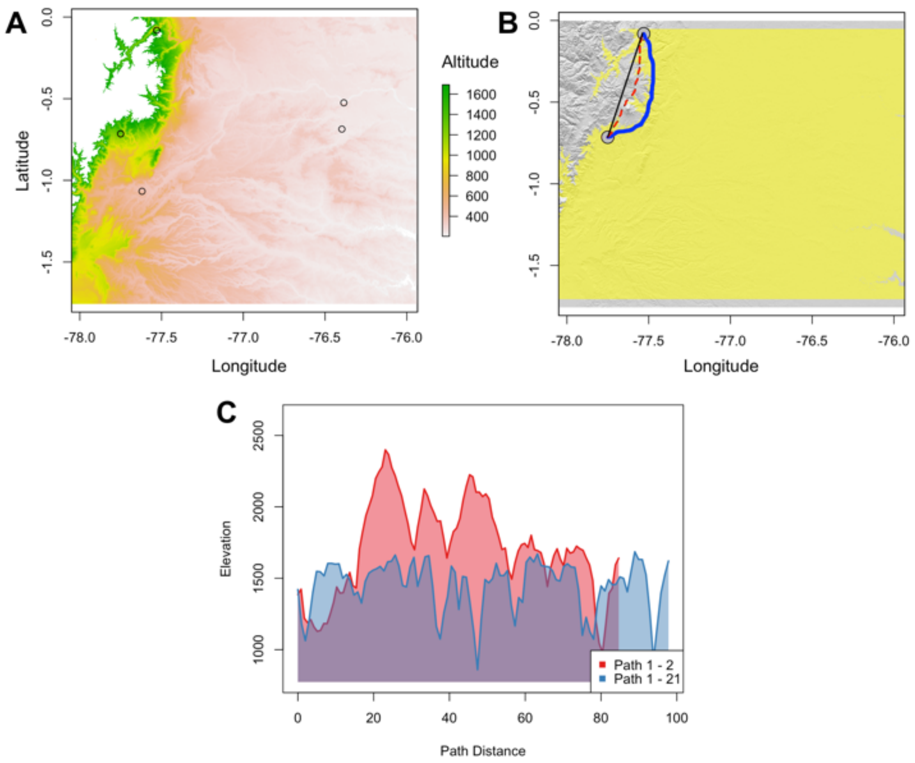


**Figure S3.** Least cost path distance example. A) Elevation raster obtained with the package elevatr [6] for highland and lowland Ecuador eastern populations (circles) with altitudes covering the elevational ranges of *H. erato*. B) Geographic distance as a straight line between populations (black line), topographic distance without accounting for elevational ranges (red dotted line), and least cost path topographic distance accounting for species elevational rages (blue solid line). The Sumaco Volcano (Napo, Ecuador) is located between these points. C) Comparison of topographic distance (red) and least cost path topographic distance (blue).

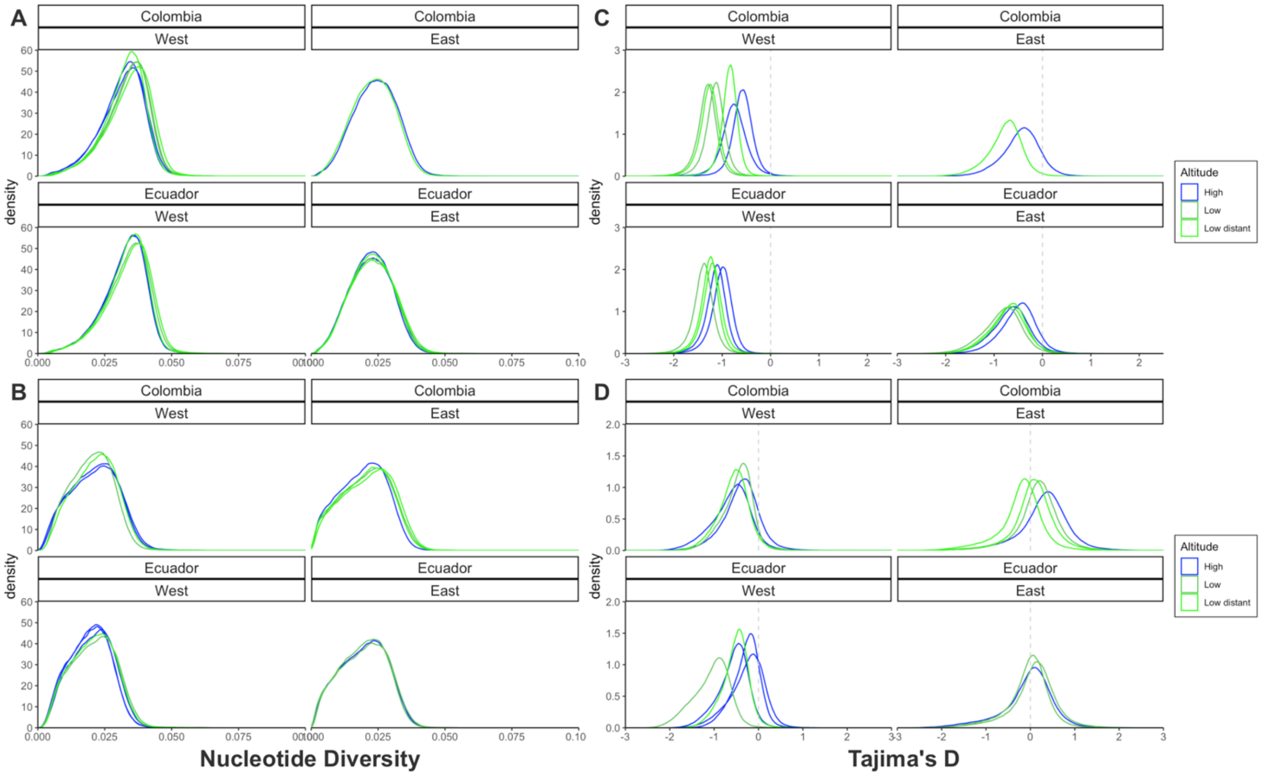


**Figure S4.** Nucleotide diversity (π, A, B) and Tajima's D (C, D) for *H. erato* (A, C) and melpomene (B, D) calculated per population (Table S1), and coloured by population area (highland, lowland, lowland distant) and by side of the Andes / species.

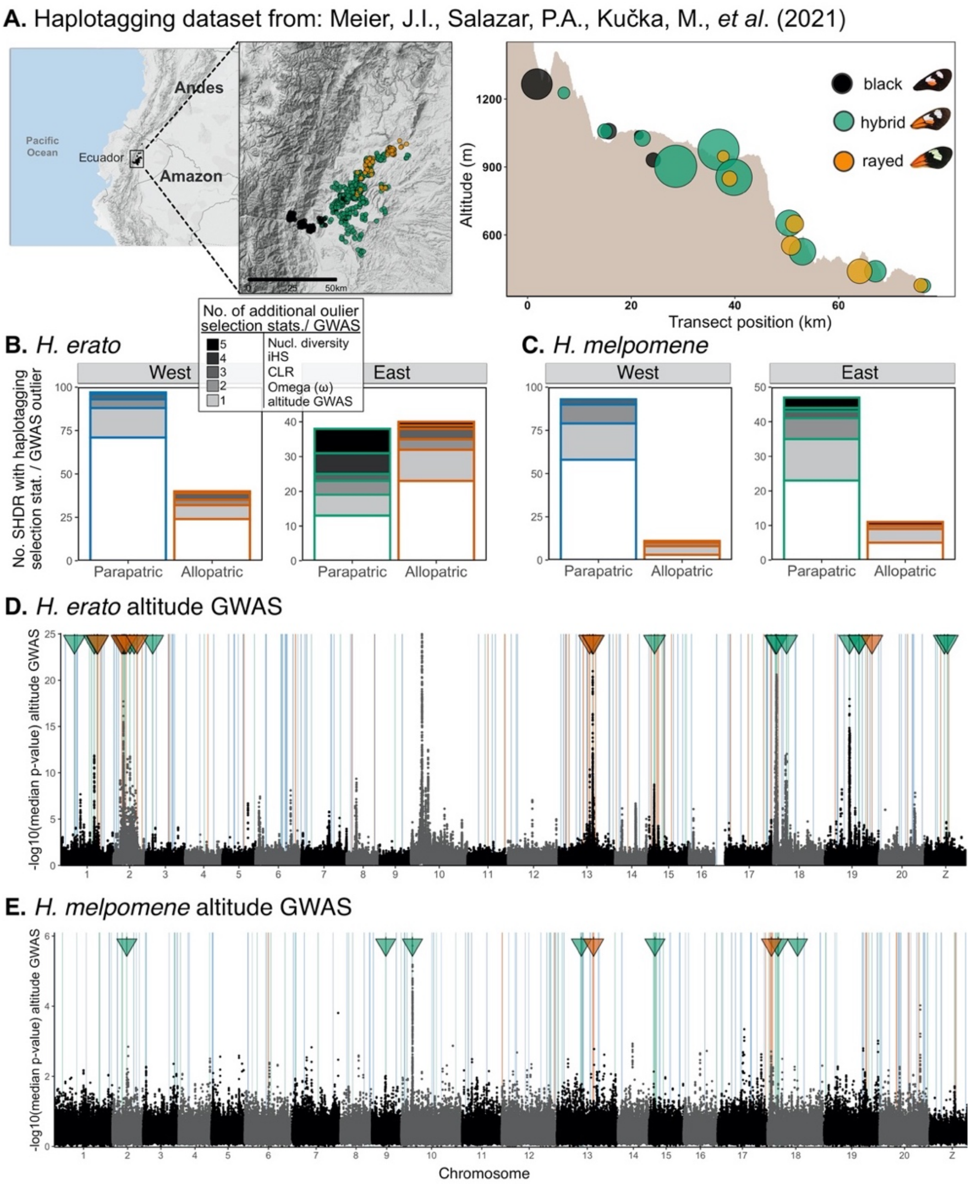


**Figure S5. Validation of SHDRs with independent published haplotagging sequencing dataset from an altitudinal cline on the Eastern Ecuadorian Andes.** A) Each point represents an individual butterfly collected in the wild and sequenced in Meier et al., 2021). The panel on the right shows the distribution of individuals sequenced across elevations. *H. erato* and H. melpomene co-occur and have three main colour pattern morphs along this cline: two distinct colour pattern morphs (*H. e notabilis* and *H. m. plesseni*, referred to as "black", and *H. e lativitta* and *H. m. malletti*, referred to as "rayed") and within-species hybrids displaying admixed phenotypes (green circles), the most common hybrid phenotype is depicted. Adapted from Montejo-Kovacevich *et al.* 2021. B-C) Number of altitude SHDR which overlap with selection statistic outliers or genome-wide association regions obtained from the analysis of the published haplotagging dataset. Darker categories indicate more selection statistics overlap (white=0 additional haplotagging selection statistics, black= 5 outlier statistics overlap). Selection statistics included: integrated haplotype score (iHS), nucleotide diversity ($\Pi$) across elevations ($\Delta\Pi = \Pi_{high} - \Pi_{low}$), omega (ω), composite likelihood ratio to detect selective sweeps using sweeD, and altitude GWAS regions (Meier *et al.* 2021). D-E) Altitude genome-wide association study of haplotagging dataset with parapatric eastern (green), parapatric western (blue), or allopatric (red) SHDRs highlighted as vertical lines. The genome-wide $90^{th}$ quantile threshold obtained for permutations was -log10(median p-value)=2.9 in *H. erato*, and -log10(median p-value)=1.8 in melpomene. Arrows indicate SHDRs that overlap with outlier altitude-associated regions.

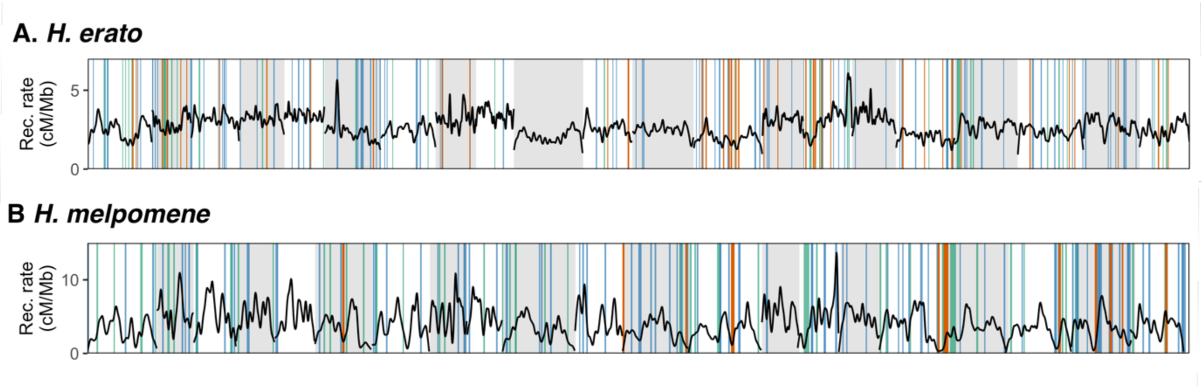


**Figure S6.** Recombination rates (cM/Mb) across the genome with SHDR highlighted as vertical lines in blue/green if shared across parapatric transects of the same species, i.e. within sides of the Andes (SHDR: blue=within West, green=within East) and in red those additionally shared across parapatric transects within species, i.e. shared across all four transects (also SHDR).

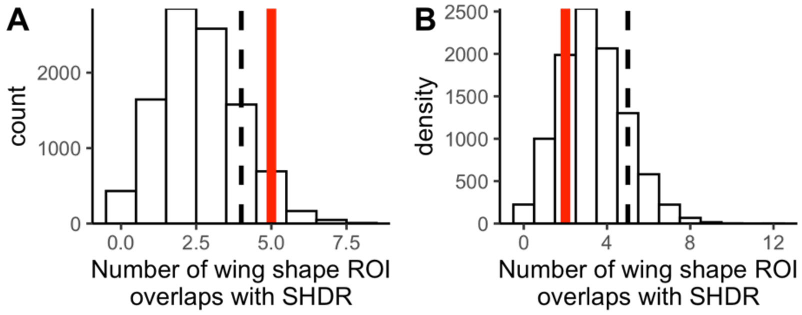


**Figure S7.** Number of wing shape regions of interest (ROI) identified in a previous study that overlap with H. erato (B) and H. melpomene (B) SHDR. Observed values are shown as red vertical lines. Bars represent 10,000 simulations, where the same number of ROI and of the same size were placed in random non-overlapping positions across the genome. Black dshed vertical line represents 90$^{th}$ quantile of simulaitons.

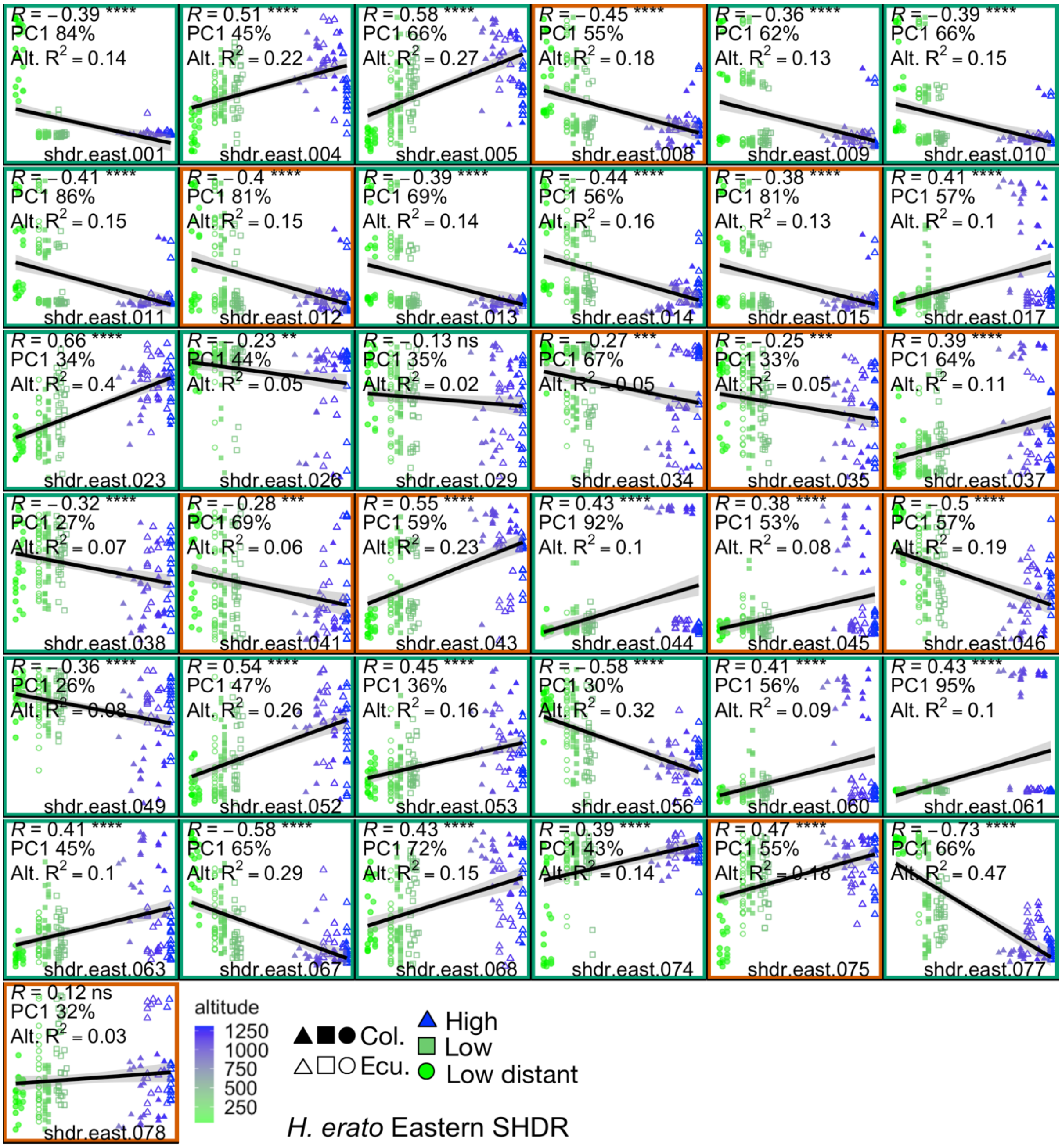


**Figure S8. Local PCA PC1 axes (y-axes) of SHDR correlate with altitude (x-axes).** *H. erato* eastern Shared High Differentiation Regions (SHDR) for which altitude was a significant predictor (P<0.05) of local PCA PC1, while accounting for neutral genome wide PCA PC1, i.e. population structure. SHDRs local PCAs included genomic windows found to be $zPBS_{high}$ outliers (>4stdv from the mean) in either Colombian or Ecuadorian clines. Each point is an individual; when solid from Colombia and empty from Ecuador. SHDRs additionally shared with the western clines (allopatric SHDRs) are highlighted with red boxes. Altitude is indicated by colour and symbols. Each SHDR plot shows: percentage of the total variation explained by PC1 (y-axes), pearson correlation coefficients (R) of the line shown (p-values for the correlations as stars ****P<0.0001, ***P<0.001, **P<0.01, *P<0.05), and the partial variation ($R^2$) explain by altitude in the model that accounts for accounting for neutral genome-wide PCA PC1. Note that *H. erato* color patter loci overlap with shdr.east.039, shdr.east.044, shdr.east.061, and the chromosome 2 inversion presented in the main text overlaps with shdr.east.008-015.

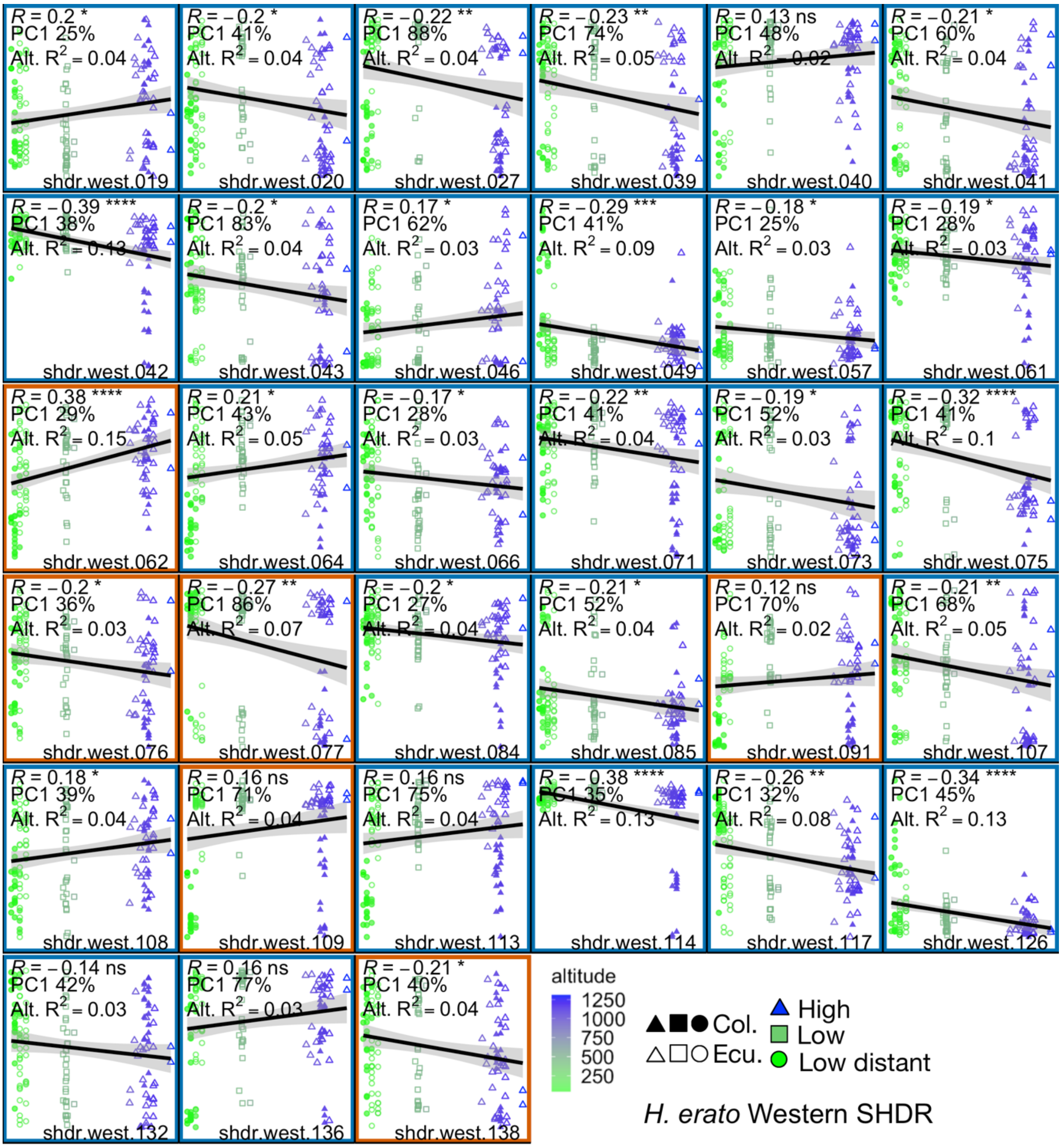


**Figure S9. Local PCA PC1 axes (y-axes) of SHDR correlate with altitude (x-axes).** *H. erato* western Shared High Differentiation Regions (SHDR) for which altitude was a significant predictor ($P<0.05$) of local PCA PC1, while accounting for neutral genome wide PCA PC1, i.e. population structure. SHDRs local PCAs included genomic windows found to be $zPBS_{high}$/zFst outliers (>4stdv from the mean) in either Colombian or Ecuadorian clines. Each point is an individual; when solid from Colombia and empty from Ecuador. SHDRs additionally shared with the western clines (allopatric SHDRs) are highlighted with red boxes. Altitude is indicated by colour and symbols. Each SHDR plot shows: percentage of the total variation explained by PC1 (y-axes), pearson correlation coefficients (R) of the line shown (p-values for the correlations as stars ****P<0.0001, ***P<0.001, **P<0.01, *P<0.05), and the partial variation ($R^2$) explain by altitude in the model that accounts for accounting for neutral genome-wide PCA PC1.

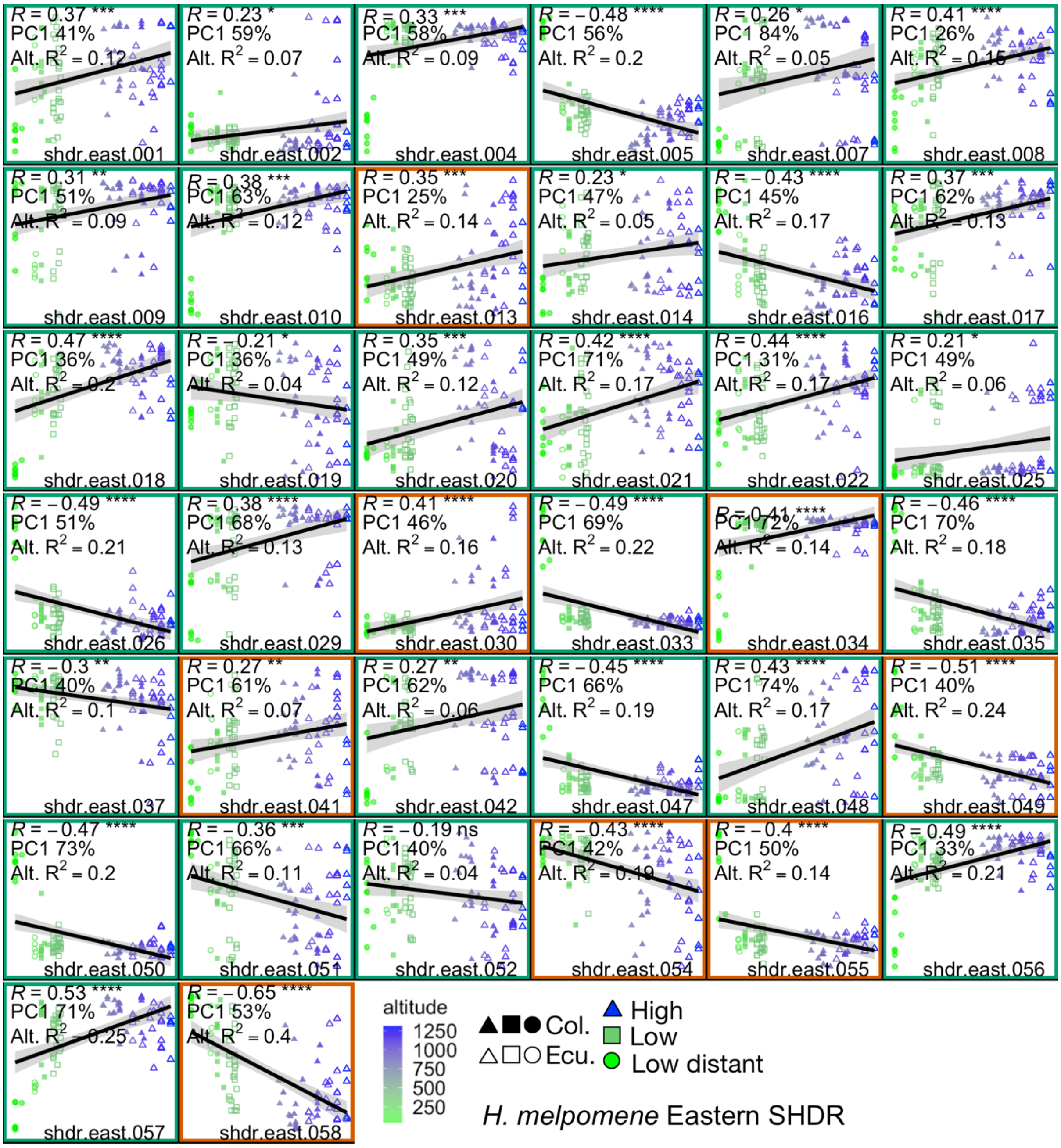


**Figure S10. Local PCA PC1 axes (y-axes) of SHDR correlate with altitude (x-axes).** *H. melpomene* eastern Shared High Differentiation Regions (SHDR) for which altitude was a significant predictor (P<0.05) of local PCA PC1, while accounting for neutral genome wide PCA PC1, i.e. population structure. SHDRs local PCAs included genomic windows found to be $zPBS_{high}$/zFst outliers (>4stdv from the mean) in either Colombian or Ecuadorian clines. Each point is an individual; when solid from Colombia and empty from Ecuador. SHDRs additionally shared with the western clines (allopatric SHDRs) are highlighted with red boxes. Altitude is indicated by colour and symbols. Each SHDR plot shows: percentage of the total variation explained by PC1 (y-axes), pearson correlation coefficients (R) of the line shown (p-values for the correlations as stars ****P<0.0001, ***P<0.001, **P<0.01, *P<0.05), and the partial variation ($R^2$) explain by altitude in the model that accounts for accounting for neutral genome-wide PCA PC1. Note that melpomene color patter loci overlap with shdr.east.024, shdr.east.036, shdr.east.042, shdr.east.043.

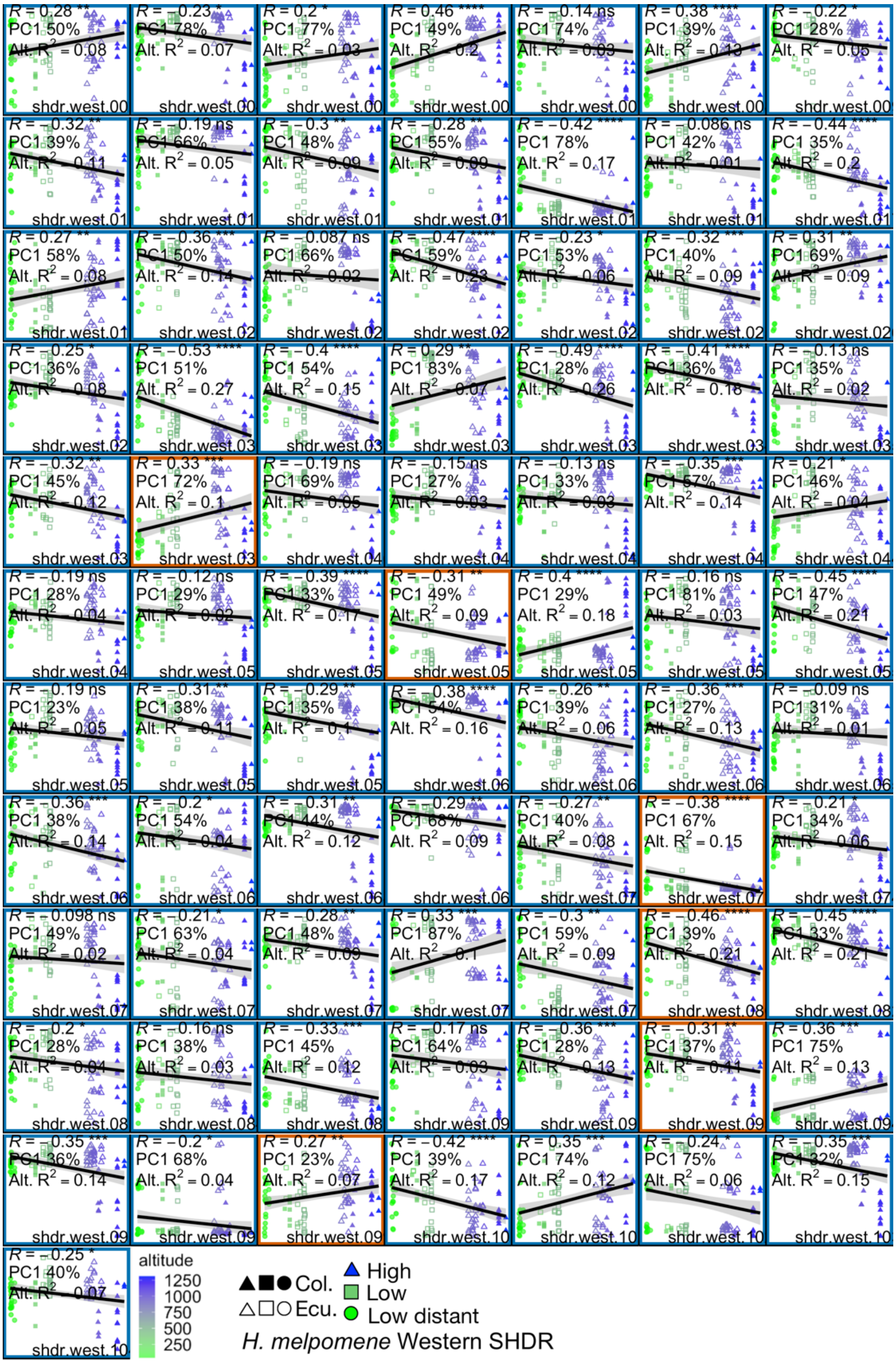


**Figure S11. Local PCA PC1 axes (y-axes) of SHDR correlate with altitude (x-axes).** *H. melpomene* western Shared High Differentiation Regions (SHDR) for which altitude was a significant predictor ($P<0.05$) of local PCA PC1, while accounting for neutral genome wide PCA PC1, i.e. population structure. SHDRs local PCAs included genomic windows found to be $zPBS_{high}$/zFst outliers (>4stdv from the mean) in either Colombian or Ecuadorian clines. Each point is an individual; when solid from Colombia and empty from Ecuador. SHDRs additionally shared with the western clines (allopatric SHDRs) are highlighted with red boxes. Each SHDR plot shows: percentage of the total variation explained by PC1 (y-axes), pearson correlation coefficients (R) of the line shown (p-values for the correlations as stars ****$P<0.0001$, ***$P<0.001$, **$P<0.01$, *$P<0.05$), and the partial variation ($R^2$) explain by altitude in the model that accounts for accounting for neutral genome-wide PCA PC1.

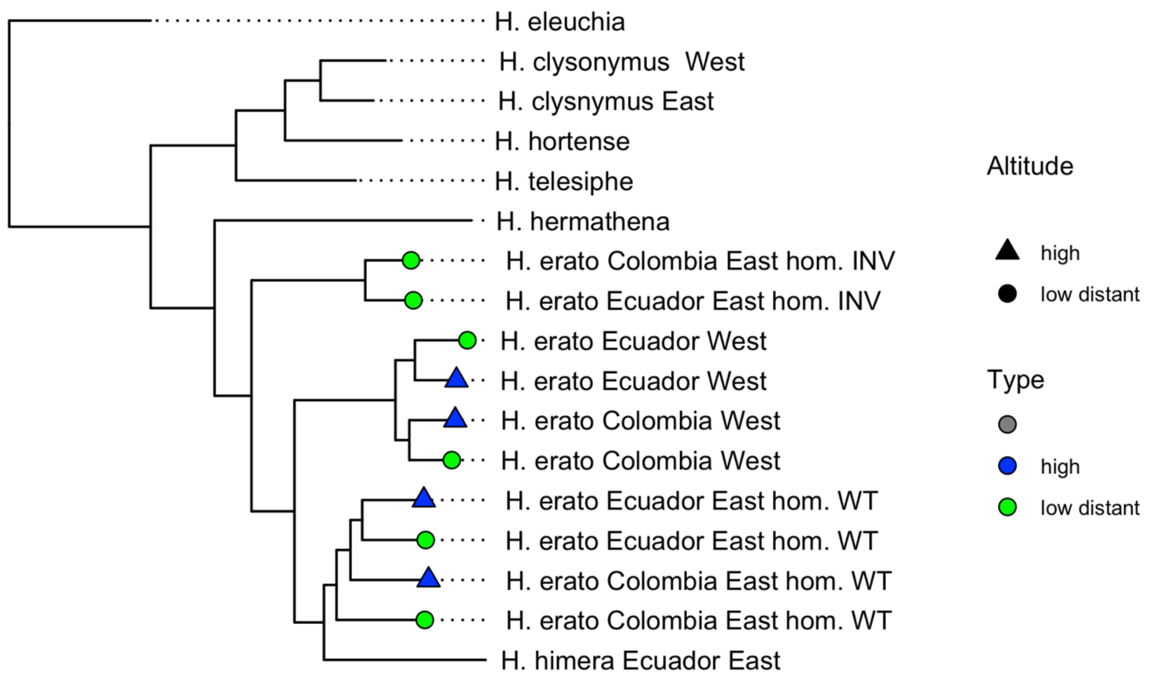


**Figure S12. Local neighbour joining tree of large putative inversion in chromosome 2 of H. erato H. erato0204:1293500 - H. erato0215:2099500.** Inversion haplotypes of eastern *H. erato* were inferred from local PCA presented in Fig 4B, INV for inverted and WT for wild-type.

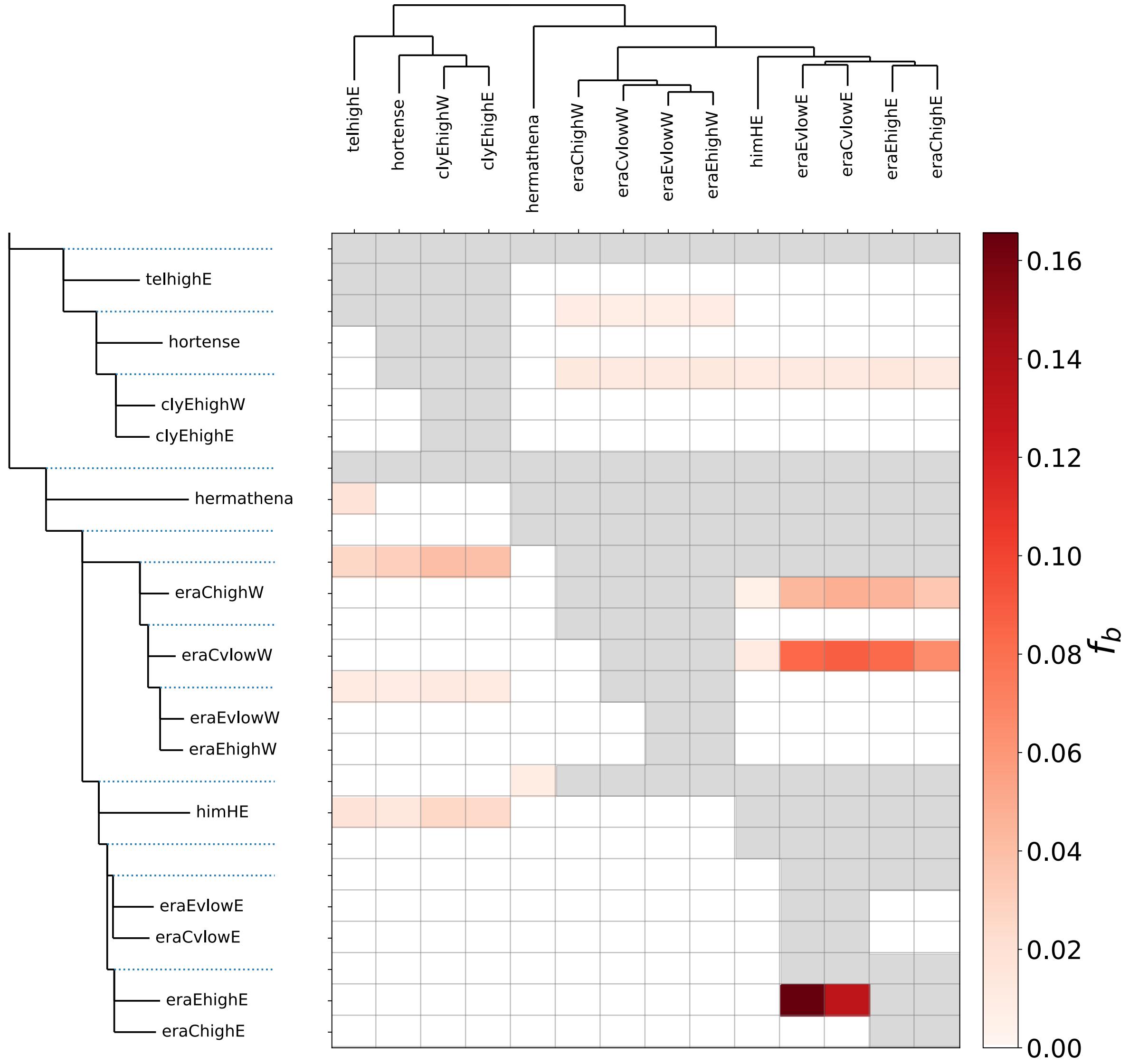


**Figure S13.** Fbranch statistics obtained with the package *DSuite* [8] for *H. erato* relatives. The 'true' tree, i.e. the most frequent tree topology genome-wide, is shown along x-axis, and is considered P3 when obtaining f-branch statistics. This tree is then expanded along the y-axis, so that each branch, including internal ones, has a row in the matrix with the inferred f-branch statistic. The values in the matrix thus refer to excess allele sharing between the branch b identified on the expanded tree on the y-axis (relative to its sister branch) and the species P3 identified on the x-axis. Non-significant f4-ratio values are set to zero prior to obtaining f-branch statistics, thus all f-branch values>0 had significant associated D statistics (p<0.001), indicating excess allele sharing for that trio.The first three letters of the tip labels correspond to species names (cly: H. clysonymus, era: *H. erato*, him: H. himera), the fourth letter, if present, indicates whether the sample was collected in Colombia (C) or Ecuador (E), then whether it was collected in the higlands (high) or lowland distant (vlow) populations, and finally the last letter indicates side of the Andes (E: east, and W: West). Outgroups are represented by the species name only.

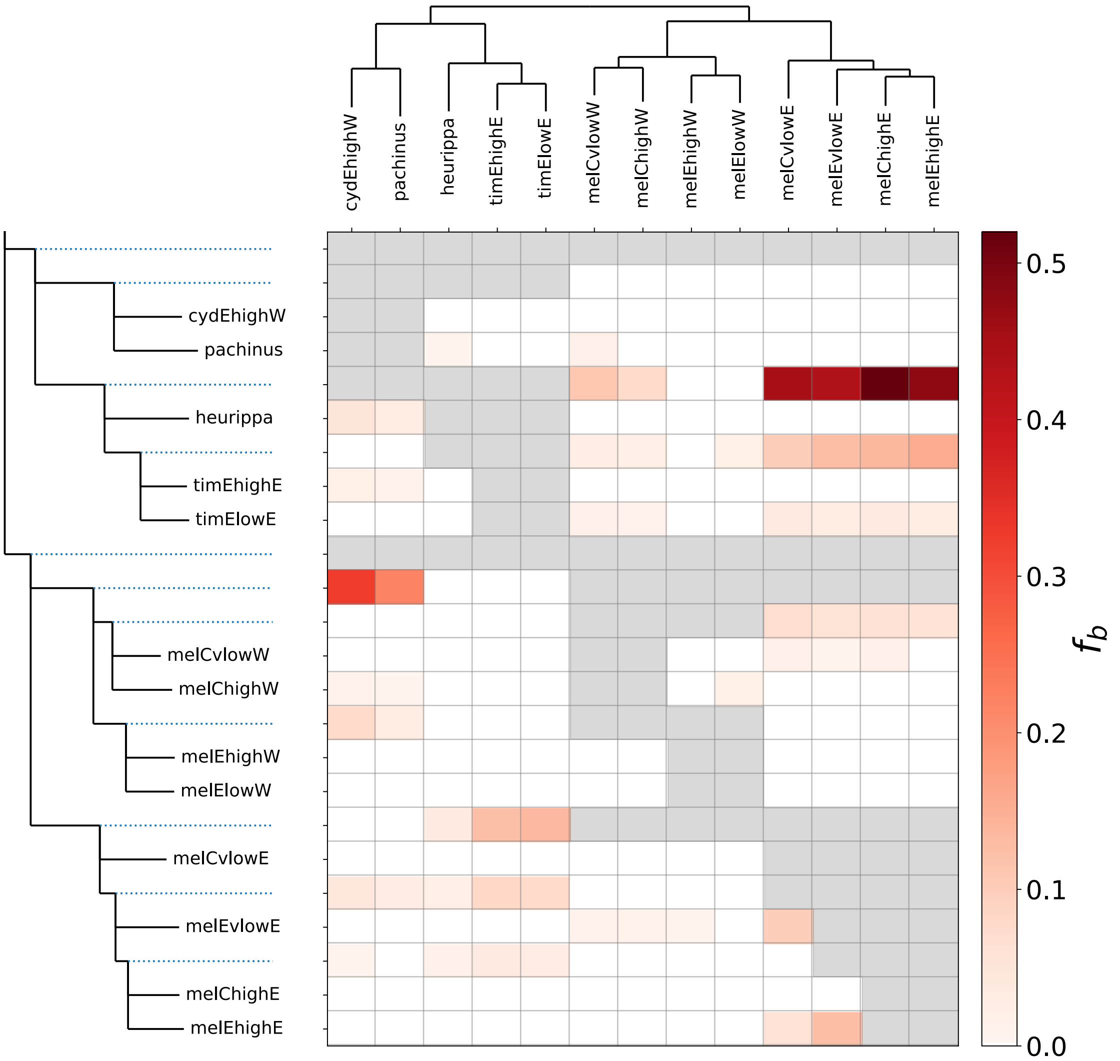


**Figure S14.** Fbranch statistics obtained with the package DSuite [8] for melpomene relatives. See explanation of axes in Figure S7. The tree was constrained to fit the species phylogeny from Kozak *et al*., 2015, to ensure that the high levels of gene flow between H. timareta and H. melpomeneThe first three letters of the tip labels correspond to species names (cyd: *H. cydno*, mel: melpomene, tim: *H. timareta*), the fourth letter, if present, indicates whether the sample was collected in Colombia (C) or Ecuador (E), then whether it was collected in the higlands (high) or lowland distant (vlow) populations, and finally the last letter indicates side of the Andes (E: east, and W: West). Outgroups are represented by the species name only.

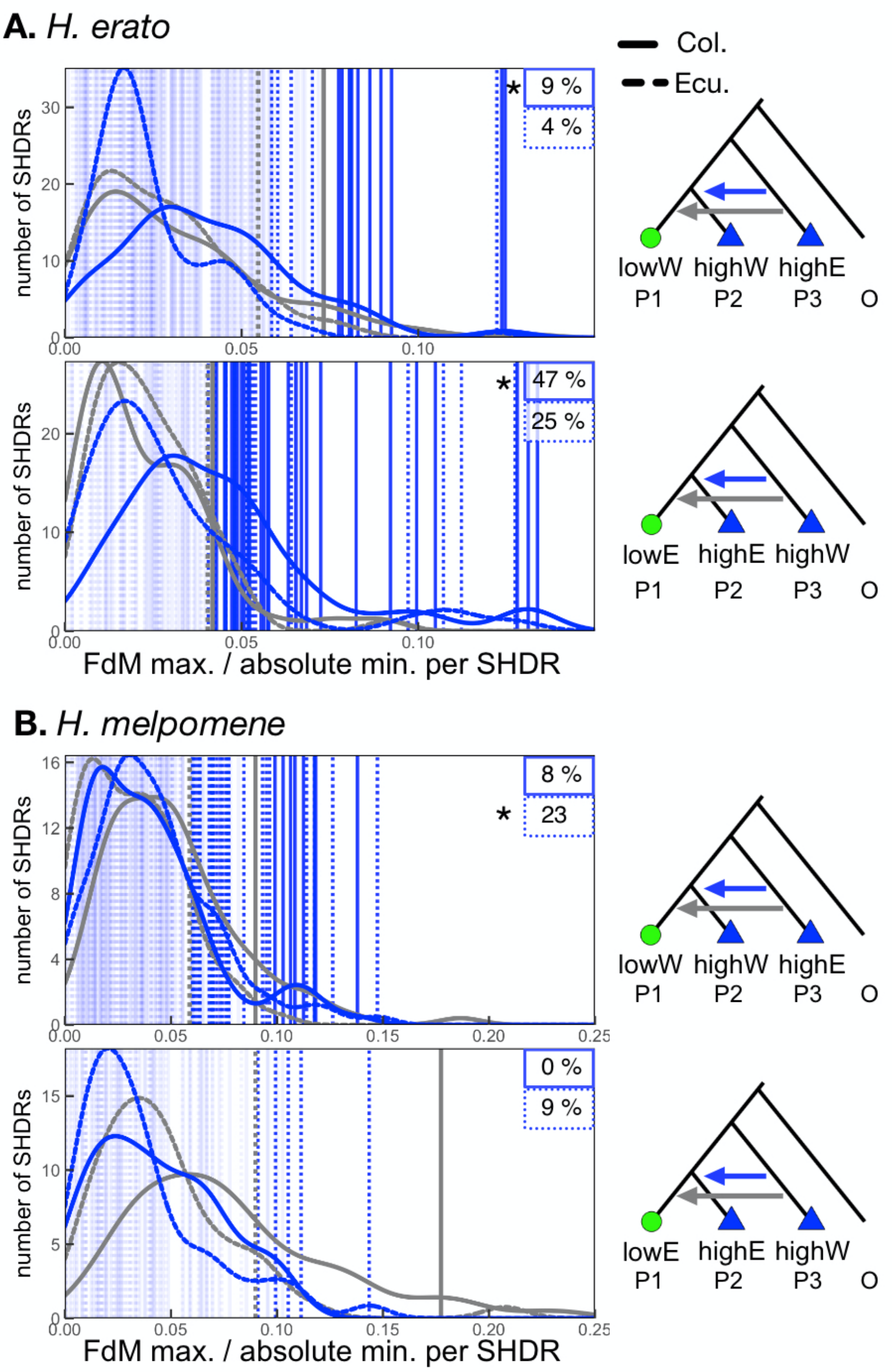


**Figure S15. Putative shared ancestral variation within high altitude shared adaptive regions (SHDRs).** Excess allele sharing between highland populations on opposite sides of the Andes, for *H. erato* (B) or *H. melpomene* (D) populations (FdM>0; blue triangles in trees) or with lowland populations (FdM<0, absolute values represented with grey lines; green circles in trees). Lowland populations correspond to lowland distant populations, except in *H. melpomene* Ecuador where only lowland populations near highlands were available. Outlier maximum FdM values per SHDR are highlighted with darker vertical lines (>90th of the neutral distribution of minimum FdM values, in grey), dashed if detected in the Colombian cline or solid if in the Ecuadorian cline. Percentage of SHDRs with outlier maximum FdM values per cline and comparison is shown on the top right corners of each plot. Significant Kolmogorov-Smirnov tests comparing overall SHDR $FdM_{max}$ and absolute $FdM_{min}$ distributions are indicated by stars.

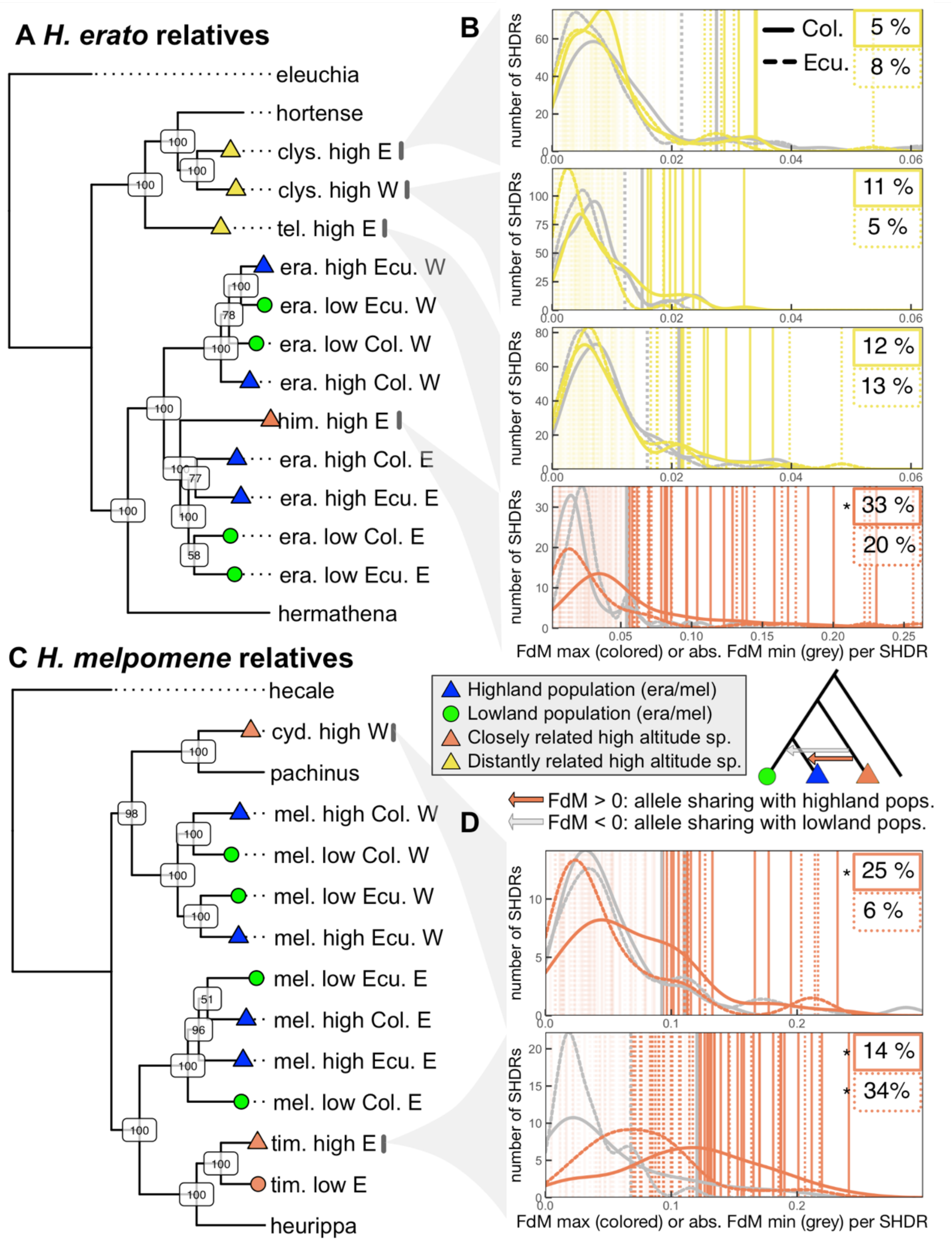


**Figure S16.** Excess allele sharing with highland *H. erato* (B) or *H. melpomene* (D) populations (FdM>0) or with lowland populations (FdM<0, absolute values represented with grey lines). Outlier maximum FdM values per SHDR are highlighted with darker vertical lines (>90$^{th}$ of the neutral distribution of minimum FdM values, in grey), solid if detected in the Colombian cline or dashed if in the Ecuadorian cline. Percentage of SHDRs with outlier maximum FdM values per cline and comparison is shown on the top right corners of each plot. Donor species (P3) of each plot in B and D is indicated with an arrow to the trees in A and C. Only parapatric allele sharing comparisons are shown, i.e. those where the recipient *H. erato* or *H. melpomene* populations were on the same side of the Andes as the high altitude species donor. Significant Kolmogorov-Smirnov tests comparing overall SHDR $FdM_{max}$ and absolute $FdM_{min}$ distributions are indicated by stars.

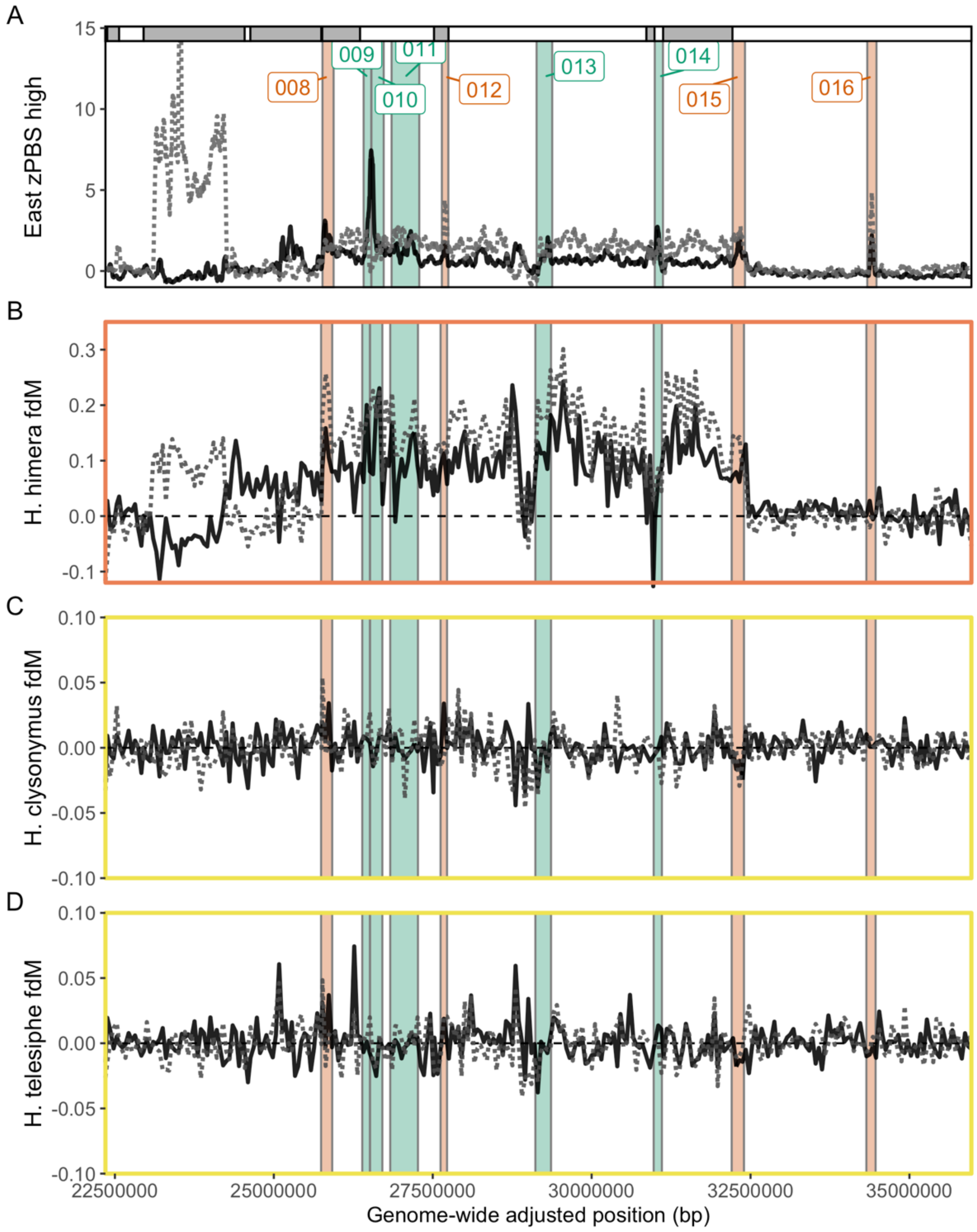


**Figure S17. H. erato chromosome 2.** A) zPBS along chromosome 2, scaffolds indicated as grey/white segments and *H. erato* Colombian cline is a dashed black line whereas Ecuador cline is grey and dashed. First elevated zPBS region, exclusive to the Ecuadorian cline, is a confirmed inversion identified in Meier et al. 2021. Second elevated region and putative inversion is presented in the main text. Excess allele sharing between highland relatives (P3: A H. himera, B H. clysonymus, C H. telesiphe) and highland *H. erato* eastern populations (FdM>0) or with lowland populations (FdM<0) in *H. erato* chromosome 2. Solid black lines represent comparisons that include the *H. erato* Colombian cline or dashed if in the Ecuadorian cline. Shared High Differentitation Regions (SHDRs) are shown as numbered vertical segments, colored by whether they represent parapatric SHDRs within Eastern clines (green) or allopatric shared across all clines (red, SHDRs shown in Fig. 2). The orange box around A denotes that himera is a closely-related species of *H. erato*, whereas in yellow are distantly related species (see Fig. S15).

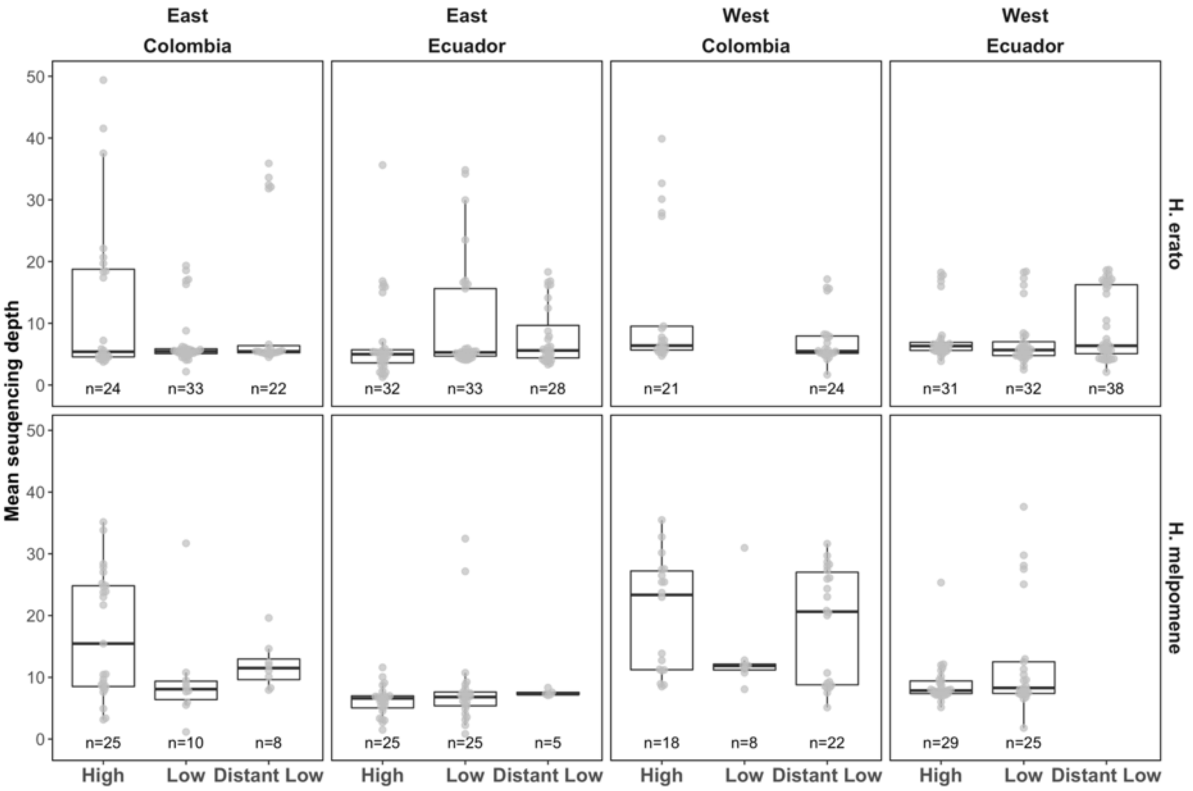


**Figure S18**. Mean sequencing depth across clines, countries, sides of the Andes and species. Genome-wide mean depth per individual calculated with samtools [9].

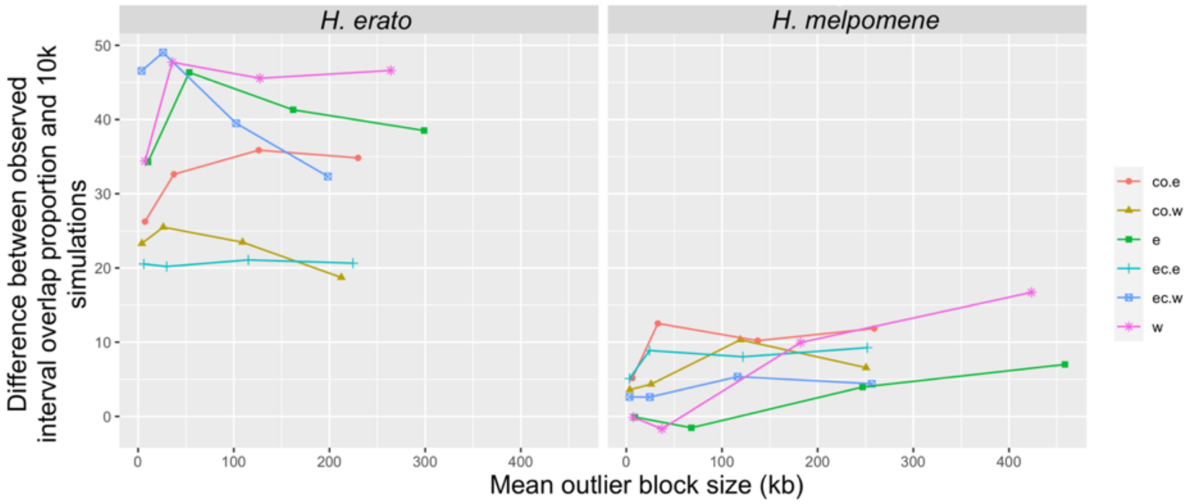


**Figure S19.** Effect of increasing outlier window buffer size on the difference between the observed HDR overlap proportion between clines and the simulated null distribution of such proportions. Each point (from left to right) represents buffers (±) of size: 0kb, 10kb, 50kb, 100kb, which results in larger HDRs (x-axis). Importantly, the difference between the observed and the simulated proportion, i.e. the significance of parallelism, only increases in *H. melpomene*, when using 100kb buffers. Each colour is a different cline population compared against the other cline on the same side of the Andes (when the code is "country.side"; co=Colombia, ec=Ecuador, e=East, w=West) or compared against the clines on the other side of the Andes (when the code is "side").
